# Diversification of the Rho transcription termination factor in bacteria

**DOI:** 10.1101/2024.06.17.599428

**Authors:** Sofia M. Moreira, Te-yuan Chyou, Joseph T. Wade, Chris M. Brown

**Affiliations:** Department of Biochemistry, University of Otago, Dunedin, Otago, 9054, New Zealand; Wadsworth Center, New York State Department of Health, Albany, NY, 12201, USA; Department of Biomedical Sciences, School of Public Health, University at Albany, Albany, NY, 12222, USA; Genetics Otago, University of Otago, Dunedin, Otago, 9054, New Zealand

## Abstract

Correct termination of transcription is essential for gene expression. In bacteria, factor-dependent termination relies on the Rho factor, that classically has three conserved domains. Some bacteria also have a functional insertion region. However, the variation in Rho structure among bacteria has not been analyzed in detail. This study determines the distribution, sequence conservation, and predicted features of Rho factors with diverse domain architectures by analyzing 2,730 bacterial genomes. About half (49.8%) of the species analyzed have the typical *Escherichia coli* like Rho while most of the other species (39.8%) have diverse, atypical forms of Rho. Besides conservation of the main domains, we describe a duplicated RNA-binding domain present in specific species and novel variations in the bicyclomycin binding pocket. The additional regions observed in Rho proteins exhibit remarkable diversity. Commonly, however, they have exceptional amino acid compositions and are predicted to be intrinsically disordered, to undergo phase separation, or have prion-like behavior. Phase separation has recently been shown to play roles in Rho function and bacterial fitness during harsh conditions in one species and this study suggests a more widespread role. In conclusion, diverse atypical Rho factors are broadly distributed among bacteria, suggesting additional cellular roles.

## INTRODUCTION

Rho is an ATP-dependent helicase that terminates transcription in bacteria by dislodging the transcription elongation complex (TEC) in a process known as Rho-dependent termination (RDT) (1). Rho primarily terminates transcription of (i) protein-coding genes in their 3’ UTRs, (ii) non-coding RNAs, and (iii) pervasive antisense transcription (2,3), thereby preventing the formation of R-loops (4).

The best biochemically characterized Rho is from *Escherichia coli* (_Ec_Rho). A monomer of _Ec_Rho has 419 amino acids (∼ 47 kDa) (Figure 1A) with three main domains - the N-terminal, RNA-binding, and ATPase domains - and it forms a ring-shaped hexamer (Figure 1B) (5–8). The _Ec_Rho N-terminal domain (Figure 1, in yellow) contains the N-terminal helix bundle (NHB) which is positively charged and facilitates interactions of the Primary Binding Site (PBS) in the RNA-binding domain (Figure 1, in pink) with specific regions of untranslated RNA known as *rut* sites (7,9). After loading onto RNA, _Ec_Rho remains as an open-ring structure (7). The ring closes when RNA engages the Secondary Binding Sites (SBS) in the ATPase domain (Figure 1, in navy blue); ring closure is also facilitated by interaction with the transcription elongation factor NusG (10). The SBS include the Q- and R-loops (Figure 1A, in green) (6,7) RNA bound to the SBS induces ATP binding and hydrolysis by the Walker A and B motifs (Figure 1A, in orange) (11), which are coordinated by the catalytic Glu, Arg valve, and Arg finger residues (Figure 1A, in blue) (12). ATP hydrolysis promotes _Ec_Rho translocation along the transcript in a 5’ to 3’ direction, until reaching the TEC and terminating transcription (11,13).

**Figure 1:**
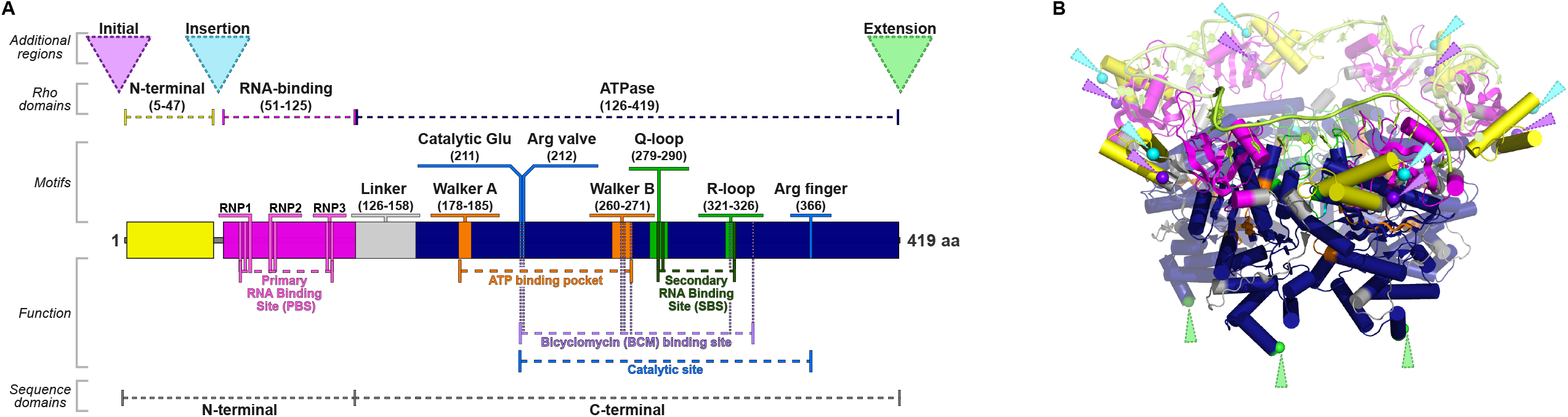
Transcription termination factor Rho of *E. coli*. (A) Diagram representing the main domains and their function within the primary sequence of Rho (B). Crystal structure of _Ec_Rho hexamer (PDB ID: 8E70). The structure is colored as shown in (B). Abbreviations: Arg, arginine; BCM, bicyclomycin; Glu, glutamate; PBS, Primary RNA-Binding Site; SBS, Secondary RNA-Binding Site.

The model of RDT described above is known as the RNA-centric model (commonly referred as the “textbook” model) and is based on extensive *in vitro* and *in vivo* characterization (13–17). This model is consistent with recent cryogenic electron microscopy (cryo-EM) structures of Rho:RNA polymerase (RNAP) complexes that also support a role for a direct Rho:RNAP interaction in the early steps of the termination process (5,18), as had been proposed previously based on biochemical data (19). Two studies described a different conformation of the Rho-RNAP complex in which Rho is inverted relative to that in the textbook model (20,21). These complexes suggest the existence of another path for Rho termination (22).

The three main domains of the _Ec_Rho sequence are relatively conserved and widely distributed, with a possibly ancient origin in Bacteria (23–25). Previous genomic analyses detected the *rho* gene in more than 90% of the analyzed bacterial genomes (706 fully sequenced genomes), with Rho absent in some small and AT-rich genomes (e.g., Mycoplasmatota, Cyanobacteriota, and certain Bacillota such as Streptococci) (23). We refer to Rho factors with only the three main domains as “typical” Rho. Atypical Rho factors with an additional region between the N-terminal and the RNA-binding domains (Figure 1, light blue dashed triangle) were frequently detected in genomes of Actinomycetota and Bacteroidota. Moreover, some species were found to contain duplicated *rho*-like genes previously classified into Group A (complete Rho with all three main domains) or Group B (incomplete Rho with missing N-terminal domain) (23).

A few atypical Rho proteins with additional regions have been shown to have additional functional properties (26–32). The atypical Rho from *Bacteroides fragilis* contains an insertion region that is essential for RDT and increases Rho’s affinity for RNA (30). *B. thetaiotaomicron* has an atypical Rho factor with an intrinsically disordered additional region that causes Rho phase separation *in vitro* and *in vivo,* promoting successful colonization in the murine gut (26). *Clostridium botulinum* has an atypical Rho with an internal prion-like region capable of self-assembling into structures resembling amyloids. The prion-like region modifies Rho conformation in response to changes in the environment (29,32). *C. difficile* has an atypical Rho with an insertion region associated with motility *in vitro* and toxin production (31). *Micrococcus luteus* has an atypical Rho with an insertion region, and this factor is capable to hydrolyze ATP in the presence of the polynucleotides poly(A) and poly(I), which significantly differs from _Ec_Rho (28). *Mycobacterium tuberculosis* has an atypical Rho with an insertion region that aids RNA binding (27).

Given the increasing number of high-quality genomes available, and the development of novel bioinformatics tools, we sought to update and extend the previous phylogenetic analysis of Rho. The distribution, phylogeny, and conservation of Rho domains were assessed among 2,730 bacterial genomes. Atypical Rho factors were detected in 39.8% of the genomes, mainly in Actinomycetota, the FCB Group (Fibrobacterota, Chlorobiota, and Bacteroidota), the PVC Group (Planctomycetota, Verrucomicrobiota, and Chlamydiota), Spirochaetota and some species of Pseudomonadota and Bacillota. Furthermore, we analyzed the sequence and predicted structure of the additional regions found in atypical Rho factors. These regions were predicted to be intrinsically disordered and to undergo phase separation, with a few being predicted to be prion-like. Our comprehensive characterization of Rho domains revealed the variation of this factor among bacteria, indicating further potential uncharacterized functions of Rho, as well as the possibility of distinct mechanisms of transcription termination occurring in species harboring atypical Rho factors. Considering these functions and the absence of Rho homologues in eukaryotes, Rho represents a promising target for the development of new antimicrobials (33). Currently, bicyclomycin (BCM) is the only approved antibiotic known to target Rho (34).

## MATERIALS AND METHODS

### Genomic data

A total of 2,784 bacterial genomes were obtained from RefSeq 204 (Table S1). These are all ‘complete’, not ‘anomalous’, ‘reference’ or ‘representative’ genome assemblies of bacterial species defined and selected by RefSeq curators to be complete representative genomes for each bacterial species (35). In addition, a small number of genomes with uncertain assignment to a clade or partial contamination with other species (n = 31) were filtered out using GUNC (Genome UNClutterer) (v1.0.4) (36) with the default thresholds (clade separation score (CSS) of > 0.45 and a contamination portion of > 2% (noted in Table S1). GUNC was designed for prokaryotes, and it has high accuracy in detecting contamination (37). A further 23 genomes were excluded as the region encoding *rho*, or the annotation of the rho genes was anomalous (Table S1, see later Methods section). The protein coding gene annotation of each genome from RefSeq was utilized (38).

### Detection of Rho domains

To gain sensitivity, the detection of Rho domains was performed by HMMER (v3.1b2) (39) with Hidden Markov Models (HMMs). The Rho HMMs from Pfam (v36.0, January 2024, Rho N-terminal: PF07498.16; Rho RNA-binding: PF07497.16; ATPase: PF00006.29) (40) were used to search against the annotated genomes using the basic command of hmmsearch. The final output was parsed with rhmmer (v0.1.0) R package (https://rdrr.io/cran/rhmmer/) and filtered based on the score cut-offs determined by Pfam (40) (Rho N-terminal: 24.9; Rho RNA-binding: 27; ATPase: 29.5).

### Construction of new specific models for Rho domains

In addition to Rho Pfam models, new specific models for Rho domains were used (Figure S1 and Figure S2). To develop these, all Rho protein sequences (COG1158) present in the eggNOG database (v5.0, March 2021) (41) (http://eggnog5.embl.de/#/app/home) were downloaded, resulting in 4,416 sequences from 3,864 species. The Pfam RNA-binding domain was detected in these sequences by hmmsearch which the matches above the score cut-off (27). Total length > 410 amino acids were selected and aligned using MAFFT (v7.394) (42) with default parameters. Multiple sequence alignment of a large dataset can be accurately and rapidly achieved with MAFFT (43). The alignments were trimmed by trimAl (v1.4.rev220) (44) with a gap threshold of 0.05, which removes gaps present in 95% or more sequences. The trimmed alignments were visualized with Jalview (v2.11.1.4) (45). Based on the 419 amino acid sequence of the Rho from *E. coli* str. K-12 substr. MG1655 (Uniprot: P0AG33), the regions corresponding to the three main domains were extracted and used to build the HMM seed models. Then, a second hmmsearch was performed using the seed models and the Pfam models against our sequence dataset to identify the coordinates of the Rho domains. After MAFFT alignments and trimmings with trimAI, the domain regions were extracted from our Rho sequences and employed to build the final Rho domain models. Next, the Rho-developed models were detected in the annotated genomes as described for the Pfam models. Besides all bacterial protein sequences in our database, we used the UniProtKB/Swiss-Prot protein sequence database (46) to test our Rho ATPase developed model.

### Comparisons of Rho in species with two *rho* genes

In species with more than one *rho* gene, we analyzed the two Rho proteins separately. Previously sequences with Rho domains had been grouped into Groups A and B (23), but no HMM models were built. However, as these groups were a mix of subclasses from different phyla, with the shorter sequence lacking the N-Terminal domain, we aligned and built new models for using the longer (A) and truncated (B) sequences. In phylogenetic analyses of species with two or more Rho proteins, we used the common RNA-binding and ATPase domains to avoid artifacts based on the presence of absence of other domains (e.g., insertion or N-terminal).

### Confirmation of species lacking Rho domains

Of the 2,753 bacterial genomes that passed GUNC analysis, 457 did not have the complete Rho (three main domains) according to the hmmsearch analysis of the translated genes from RefSeq. It is possible that a Rho ORF was interrupted or not properly annotated in some of these genomes, and thus not in the translated sequence file. To confirm this, each of the 457 genomes was completely translated in six frames using BioPerl translate to generate the amino-acid sequence of every possible protein or peptide. The hmmsearch was repeated with the Rho domain models. The Rho prediction was confirmed in 365 bacteria. The 92 divergent results were checked manually by comparing the Rho domains in closely related organisms and using tblastn to translate the genomes (47). Overall, 434 genomes did not encode complete Rho, and 23 genomes had one or more Rho domains encoded by genes unannotated by RefSeq. These were mainly short single domain ORFs and could result from a mix of genome assembly or errors, annotation errors, or possibly pseudogenes. In a conservative approach, these 23 aberrant genomes were removed from our dataset (Table S1).

### Phylogenetic analysis of bacterial species

The phylogenetic trees displaying the distribution of the Rho domains and the significant motifs were generated with PhyloT (https://phylot.biobyte.de) based on NCBI taxonomy (48). The Rho domains, genome GC%, genome size, and motifs were annotated on the trees using iTOL (v6) (https://itol.embl.de/) (49).

### Phylogenetic analysis of Rho additional regions

Additional regions between the known domains were analyzed separately. A length threshold of 25 amino acids was used as it is the minimum window size used for predictive algorithms (50). Due to the varying length of the Rho additional regions, an alignment-free method was also employed to compare these sequences. The additional protein regions were extracted, and the distance matrix in phylip format was generated with the Python script “calc_wmteric.py” from the alfpy package (51). The first line of a phylip-formatted distance matrix contains the number of sequences; the following lines have the sequence name with the pairwise distance compared to the other sequences. Next, the distance matrix was converted to MEGA format with an in-house R-script and used to construct a Neighbor-Joining tree with the MEGA (v11.0.13) software (52).

### Correlation analyses between the Rho types with bacterial traits

Correlation analyses of the Rho types/absence of Rho were performed with the following traits: (i) genetic (genome GC%, genome size, mean of the C > G bubble length in terminators, mean number of YC-hexamer repeats in terminators, number of coding genes, number of rRNA 16S genes, and number of tRNA genes); (ii) cellular (Gram stain, cell diameter, cell length, cell shape, sporulation, motility, metabolism, and doubling time); (iii) environment (isolation source, temperature range, pH range, and salinity range). Physiological and habitat information of all bacterial species of interests were sourced from the dataset provided by Madin et al. (53), which comes with NCBI species tax id. Numerical data were digitized into categorical data by quartiles.

Genomic data of bacteria species of interest, genome GC%, and genome size were sourced from NCBI RefSeq. Genomic sequences were available in FASTA format, and gene annotations were available in GFF3 format. Putative transcription terminators were identified by using the GFF3-formatted annotations which genes that are less than 55 nt apart from the neighbors are clustered as operons using the “bedtools cluster” command from BEDTools utilities (54). This criterion was based on the finding that an intergenic distance 55 nt or less predicts co-regulated genes (55). Then, putative transcription terminators were extracted as regions 250 nt downstream of all operons, where Rho-dependent terminators (RDT) were expected.

Key features of Rho-dependent terminators, namely the “C > G bubble” and the “6 x (YCN_9-13_)”, were extracted using the algorithm described by Nadiras et al. (56). Briefly, C > G bubble are regions with which the local proportion of C around each nucleotide is always greater than the local proportion of G. The 6 x (YCN_9-13_) is a series of six YC-dimers, separated by spacers of 9 to 13 nt. For each operon terminator, we computed the length of the longest C > G bubble, and the number of YC-dimer repeats with 6 YCs, and the two quantities were averaged across all operon terminators in a genome.

### Sequence alignments and logos

Protein FASTA sequences were aligned using MAFFT or CLUSTAL multiple sequence alignment by MUSCLE (v3.8) (https://www.ebi.ac.uk/Tools/msa/muscle/) (57) with the default parameters. The alignments were visualized on Geneious Prime software (v2024.0.2) (http://www.geneious.com/) and color-coded according to the polarity scheme. To represent the conservation pattern of the alignments, logos were generated with the WebLogo 3 tool (https://weblogo.threeplusone.com) (58) and color-coded based on amino acid chemistry.

### Motif discovery

Motif discovery analyses were performed with MEME (Multiple Em for Motif Elicitation) Suite (v5.5.2) (59) and MAST (v5.5.2) (Motif Alignment and Search Tool) (60) on atypical Rho sequences. First, all protein sequences of the additional regions from the atypical Rho factors were clustered using CD-HIT (v4.7) (61) with an identity-cutoff of 0.8, to generate a non-redundant FASTA file. Then, MEME was employed to detect motifs (4-40 amino acids) using the protein sequences of typical Rho factors as the background model. The discovered motifs were confirmed, and their locations on the atypical Rho sequences were annotated with MAST.

### Predictions of protein properties

The probabilities of disorder, phase separation, and prion-likeness were predicted in a representative set of Rho sequences, and the mean of scores was calculated in a 25-residue window for each Rho region in order to detect functional patches. The representative set of Rho sequences was created after clustering by CD-HIT with an identity-cutoff of 0.8. The disorder analysis and prediction of the putative function of the Rho additional regions were conducted with flDPnn (putative function- and linker-based Disorder Prediction using deep neural network) (http://biomine.cs.vcu.edu/servers/flDPnn/) (62). Spontaneous liquid–liquid phase separation and droplet-promoting regions were predicted using FuzDrop (https://fuzdrop.bio.unipd.it/) (63). Prion-like regions were predicted using PLAAC (Prion-Like Amino Acid Composition) (http://plaac.wi.mit.edu) (64).

### Tertiary and quaternary structure predictions

Predictions of monomers of Rho were retrieved from the AlphaFold Protein Structure Database (https://alphafold.ebi.ac.uk/) (65). The PDB files were downloaded, and the domains were colored on PyMOL (v2.5.2) (Schrödinger, LLC). For variants with changes in the predicted BCM-binding pocket, BCM was added with Alphafill (66) and Arpeggio (67).

### Statistical analysis

Negative and positive associations between each type of Rho and each category were tested by one-sided Fisher’s exact test. The p-value (0.001) was adjusted by Benjamini and Hochberg correction for multiple testing. All statistical pairwise comparisons were conducted via two-tailed Wilcoxon rank-sum tests.

## RESULTS AND DISCUSSION

### Bioinformatics tools and data available

The last large-scale phylogenetic analysis of *rho* was 2013 (23). Since then, many new resources and tools have become available (35,68). Additionally, the bacterial phylum names have been changed by the International Code of Nomenclature of Prokaryotes with the addition of the suffix -*ota* (69). Hence, we chose to reanalyze the phylogenetic distribution of *rho* using the bioinformatics tools and data available.

### A representative genome dataset

Previous studies have reported the presence of atypical *rho* genes, or the absence of *rho* genes in the bacterial genomes available at the time (23–25). To construct a new, high-quality representative genome dataset, we selected filtered reference genome assemblies from the RefSeq database (n = 2,784) (35). We further checked and excluded genomes with genomic chimerism, and contamination identified by the GUNC tool (31 genomes) and genomes where the *rho* gene was anomalously annotated (23 genomes, Methods). Our genome dataset comprised 2,730 high-quality bacterial genomes included representative species of all main phyla. Further details regarding the genome dataset and the hmmsearch output can be found in Table S1 and Table S2. The resulting dataset thus contains the best available genome for each species.

### Sensitivity and specificity of models for the three known Rho domains

Established models corresponding to the three classical Rho domains from Pfam were available. We also built new models including greater diversity but avoiding false positives for identifying Rho regions. The models for the “Rho N-terminal” (PF07498.16) and “Rho RNA-binding” (PF07497.16) domains exhibited good specificity, as did our new model for the RNA-binding domain (Table 1). Since both Pfam model and the new model developed here for the RNA-binding domain yielded the same results and similar sequence characteristics (Figure S1), we opted to use the established Pfam model in our analysis. In contrast, our developed model for the Rho N-terminal domain was more specific, although showing similar conserved blocks in the sequence logos (Figure S1). Given the lack of information in the literature about this domain, we also decided to use the established Pfam model for the N-terminal domain to maintain sensitivity, while potentially including some false positives.

**Table 1:** Comparison between Pfam and new models for the detection of Rho domains in bacterial annotated genomes.

| Rho domain | Number of genomes |  |  |
| --- | --- | --- | --- |
|  | Pfam model | Developed model | Pfam + new model |
| Rho N-terminal | <u>2,379</u> | 1,184 | 1,184 |
| RNA-binding | <u>2,248</u> | 2,248 | 2,248 |
| Rho ATPase | 2,730 | 2,446 | <u>2,446</u> |

The genome dataset contained 2,730 species. The numbers for the models used are underlined.

The “ATPase domain” model from Pfam (PF00006.29) is a general ATPase model and detected proteins in all bacterial species analyzed in this study (Table 1). In species where only the ATPase domain of Rho was found, it was difficult to determine if this match corresponds to the Rho ATPase domain, or another ATPase. This distinction is particularly important for species that may lack functional Rho (23). To address this issue, specific models for the Rho ATPase domain were generated. Relevant extracted regions of Rho proteins from the eggNOG database were used. Our Rho ATPase domain model detected the conserved functional ATPase and BCM-binding residues of Rho as those reported by D’Heygère et al. (23), which were not possible to distinguish using the general Pfam ATPase model (Figure S2). Indeed, functional Rho sequences had high scores (>200) for the Rho ATPase domain model and lower scores (43.8 to 113.8) for the Pfam ATPase model. In contrast, non-Rho sequences from our database and the UniProtKB/Swiss-Prot database had low scores (< 104.3) for our developed model and high scores (up to 320) for the Pfam model (Figure S3). These results confirmed the specificity and sensitivity of our “Rho ATPase” model. The hmmsearch analysis with the Rho ATPase model detected the Rho ATPase domain in 2,446 bacterial genomes (89.6%).

### Rho domain detection across bacterial species

Of the 2,730 bacterial species analyzed, the hmmsearch analysis with the “Rho N-terminal”, “Rho RNA-binding”, and “Rho ATPase” models detected at least one of the three main Rho domains in in 3,051 protein sequences from 2,499 genomes (91.5%) (Tables S2 and S3). Of these 2,499 genomes, 2,037 have only one protein sequence with Rho domains, and 462 species have multiple protein sequences with Rho domains. We define functional Rho proteins as those that have both the Rho RNA-binding and ATPase domains (Table S2). By contrast, we define “Rho-like” sequences as having only one Rho domain, or the N-terminal and the RNA-binding domains. Rho-like sequences were detected in 421 species (Table S3). Here, we focus on functional Rho factors; Rho-like sequences are described in the Supplementary Material.

### Species lacking functional Rho

In our study, 284 species (10.4%) lacked a functional Rho (Table 2 and Table S4). Species lacking functional Rho included two species of Actinomycetota (*Baekduia soli* and *Conexibacter woesei*), all members of Mycoplasmota (n = 100) and Cyanobacteriota (n = 33), 148 species of Bacillota (30.4%), notably *Streptococci* (n = 37), and a Myxococcota (*Haliangium ochraceum*) (Table 2). Details about the species lacking functional Rho are described in the Supplementary Material.

**Table 2:** Bacteria lacking functional Rho.

| Bacterial phylum | Total number of species analyzed | Number of species lacking functional Rho | Proportion |
| --- | --- | --- | --- |
| Actinomycetota | 464 | 2 | 0.4% |
| Bacillota | 487 | 148 | 30.4% |
| Cyanobacteriota | 33 | 33 | 100% |
| Mycoplasmatota | 100 | 100 | 100% |
| Myxococcota | 12 | 1 | 8.3% |

The reasons behind the absence of Rho in specific bacterial clades remain unknown. It is plausible that Rho had an ancient origin but was lost during the evolution of some lineages. Indeed, Rho is not essential in all bacteria (25,70). Therefore, the functions of Rho may be replaced by other proteins or termination mechanisms. Mycoplasmota, Cyanobacteriota, and a subset of Bacillota must use alternative mechanisms for transcription termination. Apart from Rho-dependent termination, the second well-established mechanism of transcription termination in bacteria is intrinsic termination. This factor-independent mechanism is characterized by a GC-rich structure that folds into an energetically stable stem-loop immediately upstream of a U-rich sequence (U-tract) (71). Intrinsic terminators have been reported to be abundant in bacteria devoid of Rho based on RNA-seq coupled with bioinformatic predictions. For example, putative intrinsic terminators are present in 76% of the transcripts of *Mycoplasma hyopneumoniae* (Mycoplasmota) (72). Within Cyanobacteriota, all identified transcription termination sites were classified as intrinsic terminators in *Synechocystis* sp. PCC 7338 (73) and in *S. elongatus* PCC 7942 (74). The subset of Bacillota lacking Rho includes the *Streptococcus* species, which also exhibit a notably high proportion of transcripts with intrinsic terminators as reported for *S. agalactiae* NEM316 (92%) (75) and *S. pyogenes* S119 (89%) (76).

### Classification of Rho into four major structural types

In _Ec_Rho (419 amino acids), the N-terminal domain is located near the beginning of the sequence (starting at the fifth residue) and is separated from the RNA-binding domain by four residues (EDIF); the ATPase domain has 293 residues (Figure 1A). However, many sequences of functional Rho analyzed here (n = 1146) are longer than _Ec_Rho, due to additional regions located before, between, or after the typical Rho domains (Figure S4A).

We classified additional regions based on their position in the Rho sequences and named Rho additional regions over 25 amino acids long as follows: (i) “initial region”: located before the N-terminal domain, (ii) “insertion region: located between the N-terminal and RNA-binding domains, (iii) “extension region”: located after the end of the Rho ATPase domain. An > 25 amino acid additional region between the RNA-binding domain and the Rho ATPase domain was not observed. Some sequences lacking the N-terminal domain had an additional region before the RNA-binding domain which we called the “2d” region. The 2d region may correspond to a remnant left after the loss of a detectable N-terminal domain or might functionally replace the N-terminal domain (analyzed below).

We define four major structural types of functional Rho factors, which we call Type 1 (typical Rho with the three main domains), Type 2 (atypical Rho with an initial and/or an insertion region), Type 3 (atypical Rho with an extension region), Type 4 (atypical Rho lacking the N-terminal domain) (Figure 2A). A single category for Rho with initial or insertion regions was chosen based on the structure of Rho (Figure 1B; PDB ID: 8E70), which indicates the likely proximity of these two additional regions. Four subtypes of the Rho Type 2 (2a, 2b, 2c, and 2d) with all the possible positions of the initial, insertion and 2d regions were also possible in our classification (Figure S4B; described in detail in the Supplementary Material). The main classification in four types captures the structural variability in Rho that likely contributes to its function. Type 2 extends previous ideas of insertion regions, and Type 4 refines the previous idea of a shorter “group B” Rho (23).

**Figure 2:**
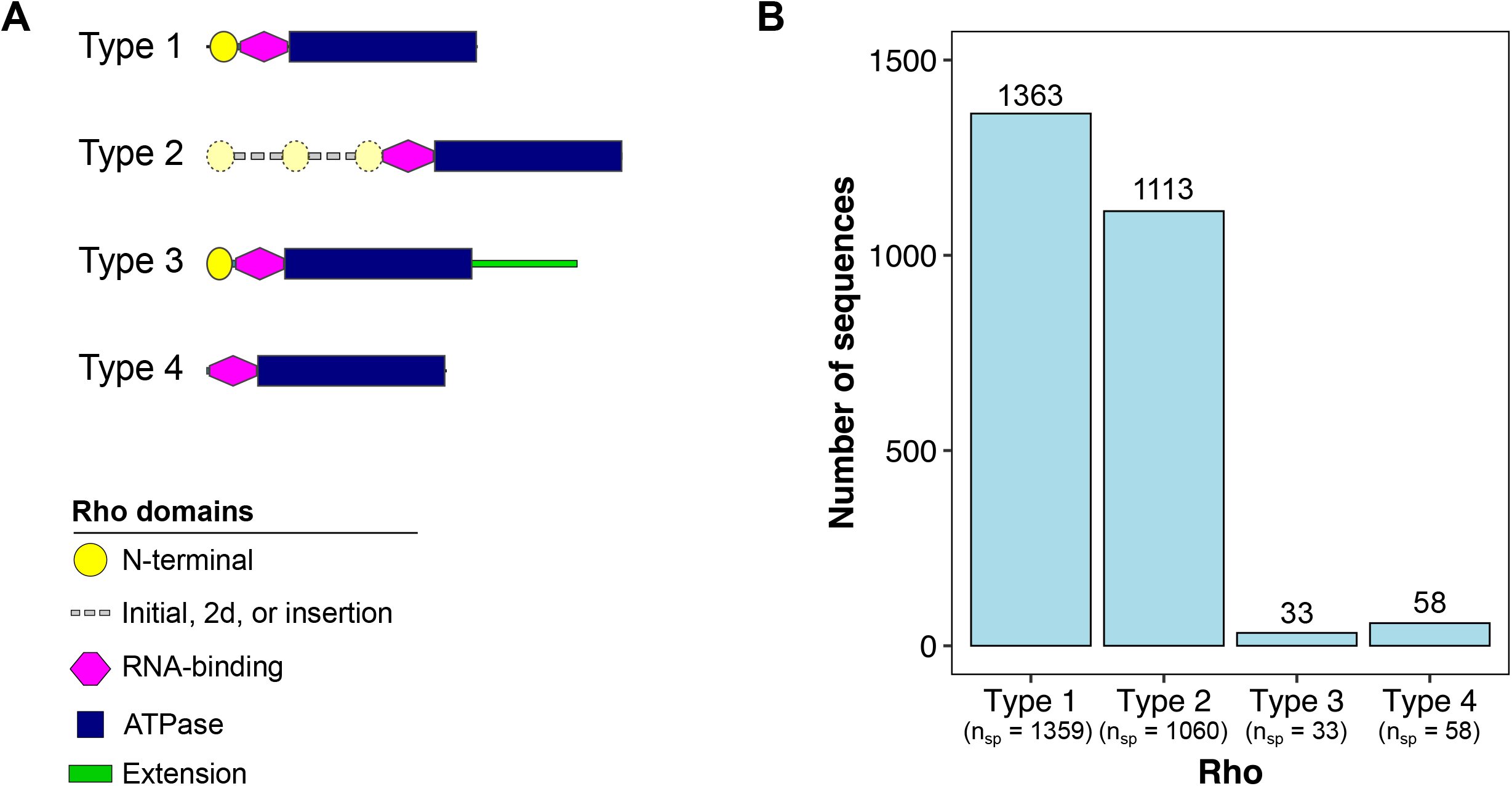
Four major types of functional Rho. (A) Schematic representation of the four Rho types. Type 1 (typical Rho with the three main domains), Type 2 (atypical Rho with an initial and/or an insertion region), Type 3 (atypical Rho with an extension region), Type 4 (atypical Rho lacking the N-terminal domain). Rho Type 2 might have the N-terminal domain after the initial region (2a), at the beginning of the sequence (2b), between the initial and insertion regions (2c) or the additional region may lack a recognisable N-terminal domain (2d). More detail is in Figure S4. (B) Number of sequences of each Rho type. There are also 284 species in our dataset that lack the functional Rho. Abbreviations: n_sp_ = number of species.

Functional Rho proteins are mainly Type 1 (n = 1363) and Type 2 (n = 1113) (Figure 2B). Type 3 had only 33 sequences while Type 4 had 58 sequences (Figure 2B). More than one copy of Rho (usually two) was observed in 116 bacterial genomes, explaining the difference between the number of species and the number of sequences. The distribution of the Rho types according to phylogeny and bacterial characteristics are presented below.

The essential role of the three main domains in _Ec_Rho has been extensively confirmed through mutagenesis (16,19,77–79). Each of the distinct types of Rho proteins has been shown to be or is likely to be functional. Indeed, the termination activity of Rho with different atypical Type 2 sequences has been reported, such as Rho Type 2a of *Xanthomonas campestris* (80,81) and *X. oryzae* (80), and Rho Type 2b from *B. fragilis* (30), *B. thetaiotaomicron* (26), and *M. luteus* (28), as well as Rho Type 2c from *M. tuberculosis* (82–84).

In contrast, Rho types that lack the N-terminal domain (notably Rho Type 4) may have modified activity in initially binding RNA (9,85). Deletion of the 27 first N-terminal residues in _Ec_Rho including the seven basic residues of the N-terminal domain abolished the capacity of this factor to bind RNA (79). In *Fusobacterium nucleatum,* which has only a Rho Type 4, the *rho* gene was shown to be one of the essential genes (86), indicating the functionality of this Type.

### Rho types are diversely and widely distributed among bacterial genomes

The distribution of Rho was analyzed across the bacterial phylogenetic tree. Figure 3A depicts the distribution of the Rho types among 12 representative bacterial phyla. Figure 3B shows a phylogenetic tree based on the NCBI taxonomy (280 diverse species), with each tip representing the detected Rho sequence(s). In addition, the bacterial phyla, genome size and genome GC% are shown with different colors. Figure S5 displays the complete phylogenetic tree with all the species analyzed in this study (n = 2,730) (data in Table S2).

**Figure 3:**
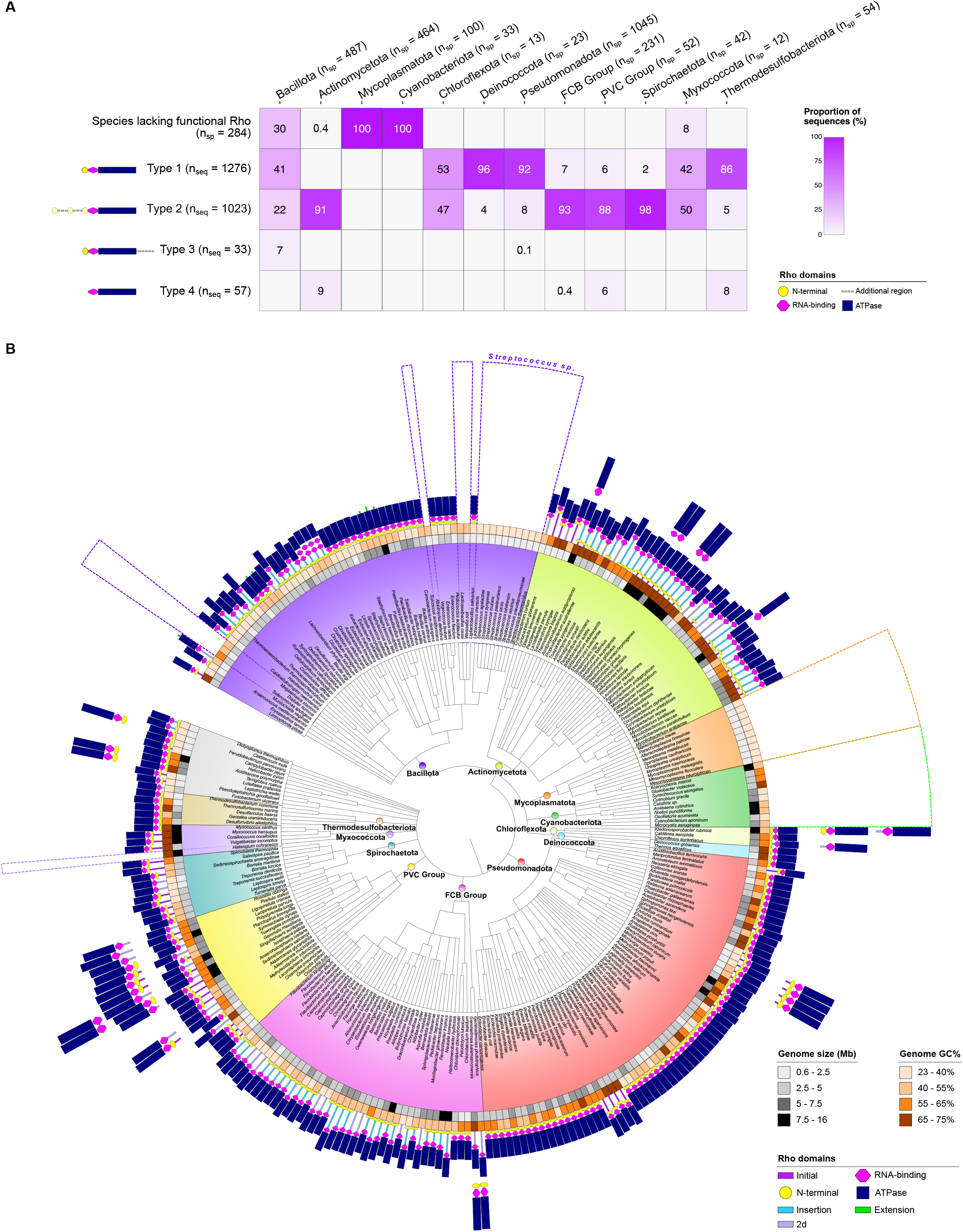
Rho factors in representative bacterial species. (A) Distribution of Rho types among bacterial phyla. Each column of the heatmap shows the proportion of species lacking Rho and the proportion of sequences of each functional Rho type in 12 representative bacterial phyla. Table S5 contains this distribution for all 37 bacterial phyla analyzed in this study. (B) Distribution of Rho types in bacterial genomes associated with a phylogenetic tree based on the NCBI taxonomy. Of the 2,730 genomes analyzed in this work, 280 species are shown in this tree to facilitate the visualization of the distinct types of Rho domain distribution (depicted as geometric forms). Dashed boxes highlight the species in which no or only the Rho N-terminal domain was detected. Genome size (Mb) and GC content (%) are also depicted. Abbreviations: n_sp_: number of species; ns_eq_: number of sequences.

At the phylum level, Bacillota has the most diverse distribution of Rho types (Figure 3B, purple). About 41% (n = 200) of Bacillota species have Rho Type 1, while 22% of the members (n = 107) have Rho Type 2 and 6.6% (n = 32) have Rho Type 3. However, 30.4% (n = 148) of Bacillota appear to have lost Rho and do not have Rho domains at all (e.g., Negativicutes, *Streptococcus*, and most members of the Lactobacillaceae) (Figure 3B, purple dotted sectors). All members of­­ the Phylum with the smallest genomes (<2 Mb) Mycoplasmatota (n = 100) lacked Rho (Figure 3B, orange), but Cyanobacteriota with larger genomes (2.5-7.5 Mb, n = 35, Figure 3B, green) also lack functional Rho. Taken together, this likely indicates independent loss of ancestral *rho* in up to five clades (Figure 3B, dotted sectors).

In contrast, nearly all Actinomycetota (n = 460) have Rho Type 2 with insertion regions (Figure 3B, light green). The exceptions are two species (*Baekduia soli* and *Conexibacter woesei*) only having Rho-like sequences (N-terminal + RNA-binding domains) and other two species (*Streptomyces cyaneogriseus* subsp. *noncyanogenus* and *Streptomyces hundungensis*) having Rho Type 4.

In Pseudomonadota (n = 1045), most species (92.5%) have Rho Type 1, like *E. coli* and *Pseudomonas aeuriginosa*. A few members only have Rho Type 2 (n = 85) or Type 3 (n = 1) (Figure 3B, red).

The majority of species from the FCB Group (Fibrobacterota, Chlorobiota, and Bacteroidota) (Figure 3B, pink), PVC Group (Planctomycetota, Verrucomicrobiota, and Chlamydiota) (Figure 3B, yellow), and Spirochaetota (Figure 3B, dark green) have Rho Type 2.

Overall, the distribution of Rho types is related to the phylogeny. Species from the same genus often have the same type of Rho, with the atypical Rho factors mainly detected in Actinomycetota, the FCB and PVC Groups, Spirochaetota, and a fraction of Pseudomonadota and Bacillota.

### Species with two Rho genes

Two or more Rho genes were detected in 116 genomes. They were mainly from Actinomycetota (n = 62) and the PVC Group (n = 30) (Figure 3B, Figures S6 and S7). It had previously been noted that some species contained an additional Rho, usually lacking the N-terminal domain but collectively clustered as ‘Group B’ (23). In our dataset, ‘group B’ consists mainly of longer (Subtype 2d in Actinomycetota and the PVC Group) and shorter Rho (Type 4) (Figure S6). We built HMM models of the common region to compare the N-terminal containing (Group A) to the second, shorter Rho (Group B) (Methods).

The alignment of these pairs of sequences showed that the longer sequence (A) is very conserved and has all three conserved Rho domains. In contrast, the shorter one has an RNA-binding domain with some small insertions, but the catalytic residues in the ATPase domain and BCM-binding pocket are conserved (Figure S8).

These results suggest the second copy of Rho rarely arises by recent duplications of Rho, but also suggests ancient duplication in several clades (see also Supplementary Text). As Rho is a hexamer there is a possibility of hetero-hexamers, where the missing N-terminal domain function could be provided in trans by Type 2, which contains both additional regions and an N-terminal domain. Both the N-terminal domain and basic additional regions in this species have basic patches that could non-specifically bind RNA or melt GC rich RNA (85).

### Associations between Rho Types and bacterial traits

Besides phylogeny, we assessed the distribution of the Rho types according to genetic, cellular, and environmental bacterial available features (Figure 4, Figures S9 and S10, and Table S6). Overall, the correlation between the Rho types/absence of Rho with cell and habitat traits do not reveal additional insights beyond those found with phylogeny (Figure S9). For example, Type 2 has a significant positive association with sporulate Gram-positive bacteria isolated from soil, traits expected in Actinomycetota, the phylum with the greatest proportion of Type 2 sequences. The most significant and interesting correlations are regarding the genome (genome size and GC%) and terminator characteristics (mean of C > G bubble length and mean number of 6 x (YCN_9-13_) in terminators) (Figure 4).

**Figure 4:**
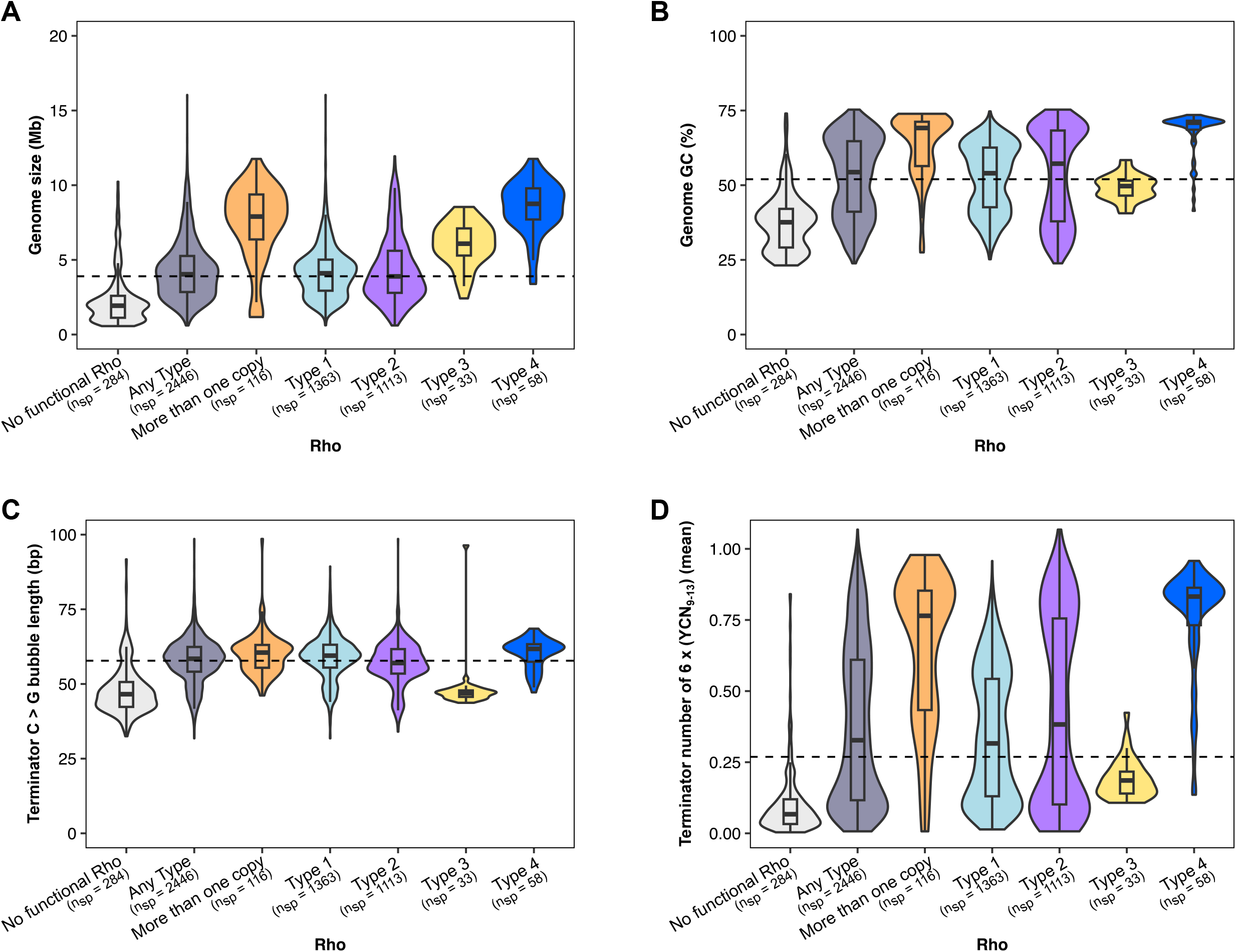
The genomic features of bacterial species containing Rho groups. (A) Genome size (Mb). (B) Genome GC (%). (C) Mean C > G bubble length in terminators (bp) (D) Mean number of 6 x (YCN_9-13_) in terminators. The dashed line indicates the median values for genome size (3.86 Mb), GC% content (52%), mean C > G bubble length (bp) (57.8 bp), and mean number of 6 x (YCN_9-13_) in terminators (0.27). Box plots depict the distribution of the values as quartiles. The line at the center of each boxplot corresponds to the median value. Violin plots represent the density distribution of the values. Pairwise p-values for the panels A-D are shown in Table S7.

Clearly, species lacking functional Rho are distinct from the other species (p < 0.001). They have significant positive associations (p < 0.001) with small low GC genomes and weak terminators (short C > G bubbles and nearly zero 6 x (YCN_9-13_)). These results are consistent with termination mechanisms that do not involve Rho. Conversely, species with any type of Rho show significant positive associations with medium to large GC rich genomes, a tendency observed before (23), but also association with long C > G bubbles and 6 x (YCN_9-13_). However, there are significant differences between and within the Rho types.

Rho Type 1 is positively associated with medium size genomes (2.5-5 Mb) of varying GC% (45-65%) and variable C > G bubble length (58-100 bp) and number of 6 x (YCN_9-13_) (0.1-0.3). This variability is due to the Bacillota and the Pseudomonadota phyla that have opposite characteristics (Figure S10). In the case of Bacillota, it is possible that RDT is not the main mechanism of transcription termination in these organisms, a fact supported by the non-essentially of *rho* in *Staphylococcus aureus* (25) and the lack of Rho in 30% of this phylum (this study).

Our correlation analyses revealed that Rho Type 2 is positively associated with large (7.5-26 Mb), high GC genomes (65-75%) with medium C > G bubble (52-58 bp) and 6 x (YCN_9-13_) (0.6-1) in terminators. In this case, an initial/insertion region might provide Rho additional properties that facilitate the process of transcription termination in those genomes (e.g., Actinomycetota and the PVC Group) by melting the RNA secondary structures, for example. However, members of Bacillota, Spirochaetota, and the FCB Group with Rho Type 2 have low GC genomes with few repeats of YCN_9-13_ in the terminators (Figure S10). In such organisms, the additional regions of the Rho Type 2 might permit distinct functions from those with high GC genomes, as seen in *B. thetaiotaomicron* (26).

A different scenario was observed for Rho Type 3. Differently from the rest of Bacillota, Rho with a C-terminal extension region was positively associated with medium values of genome size (5-7.5 Mb), GC% (40-55%) and number of 6x(YCN_9-13_) (0.1-0.3) in terminators but short C > G bubble length (30-52 bp).

In contrast, Rho Type 4 have similar results to Actinomycetota with Type 2, being positively associated with large (7.5-16 Mb), high GC% (65-75%) genomes with 6 x (YCN_9-13_) in terminators. The same results were observed with species containing more than one copy of Rho excepting *Anaplasma* sp. and *Ehrlichia* sp. (Pseudomonadota) that have exact duplicates of Rho Type 2 sequences.

### Variations within the three main Rho domains

Currently, knowledge about the essential residues in Rho associated with active transcription termination mainly comes from mutagenesis assays with _Ec_Rho (14,16,17,87). How conserved are these residues in atypical Rho sequences? To investigate this, we extracted each Rho domain detected by the hmmsearch analysis and aligned the sequences by the Rho groups, considering also the structures of Rho (PDB IDs: 8E3H, IPV0, and IPV4) (5,34,88). Sequence logos were built to visualize the most conserved residues; figures for each domain are present in the Supplementary Material (Figures S11, S12, and S15).

### The Rho N-terminal domain

The N-terminal domain of _Ec_Rho is positively charged due to the presence of a few patches of lysine and arginine residues and has been proposed to be involved in non-specific binding to the RNA backbone (9,85). Our analysis indicated that sequences of this domain had some partially conserved blocks (e.g., R(M/L)RK) (Figure S11, 27-30 in Rho Type 1). The sequences of the N-terminal domain from Rho Type 2 were more divergent, but still contained positively charged residues, suggesting that these divergent and widely distributed sequences may play the same role as the typical N-terminal domain in recruiting RNA. In fact, this domain belongs to the helix-extended loop-helix (HeH) superfamily (Pfam: CL0306), which consists of 16 nucleic acid binding folds (89). Some of this clan bind RNA e.g., SAP (SAF-A/B, Acinus and PIAS proteins) motifs (90), which have been found in RNA processing complexes in yeast being associated with degradation of apoptotic chromatin, DNA repair, and RNA processing (91). In structures of Rho complexes this domain is on the outside, with both ends near the RNA-binding domain (Figure 1B).

### The Rho RNA-binding domain

The RNA-binding domain of _Ec_Rho contains the RNA primary binding sites (RNP1, RNP2, and RNP3) as part of a cold-shock single-stranded RNA binding domain. Previous structures of _Ec_Rho revealed the residues within RNP1, RNP2, and RNP3 that bind specific pairs of bases in the *rut* sites (Figure 1A) (5,7,18,92,93). Our analysis demonstrated that the RNA-binding domain is conserved in the four types of functional Rho (Figure S12), and some exceptional cases were observed (Figure 5). Four species of Rho Type 2 from the Eubacteriaceae family (Bacillota) have duplicated, tandem RNA-binding domains (Figure 5A) separated by a linker of 20 amino acids with poor Alphafold prediction. Duplication was also seen in 23 other Eubacteriaceae from Genbank, not in our high-quality set. The first copy of the RNA-binding domain in those species contains some point substitutions compared to _Ec_Rho (Figure 5A and Figure S13). However, the RNA primary binding sites of these duplicated domains contains all the critical residues, indicating that they are likely functional and able to bind to RNA. Indeed, the Alphafold structures of the duplicated RNA-binding domains were predicted to have the same conformation as _Ec_Rho (PDB ID: 8E6W) potentially binding in the same way to the *rut* site from that complex (Figure 5B and 5C).

**Figure 5:**
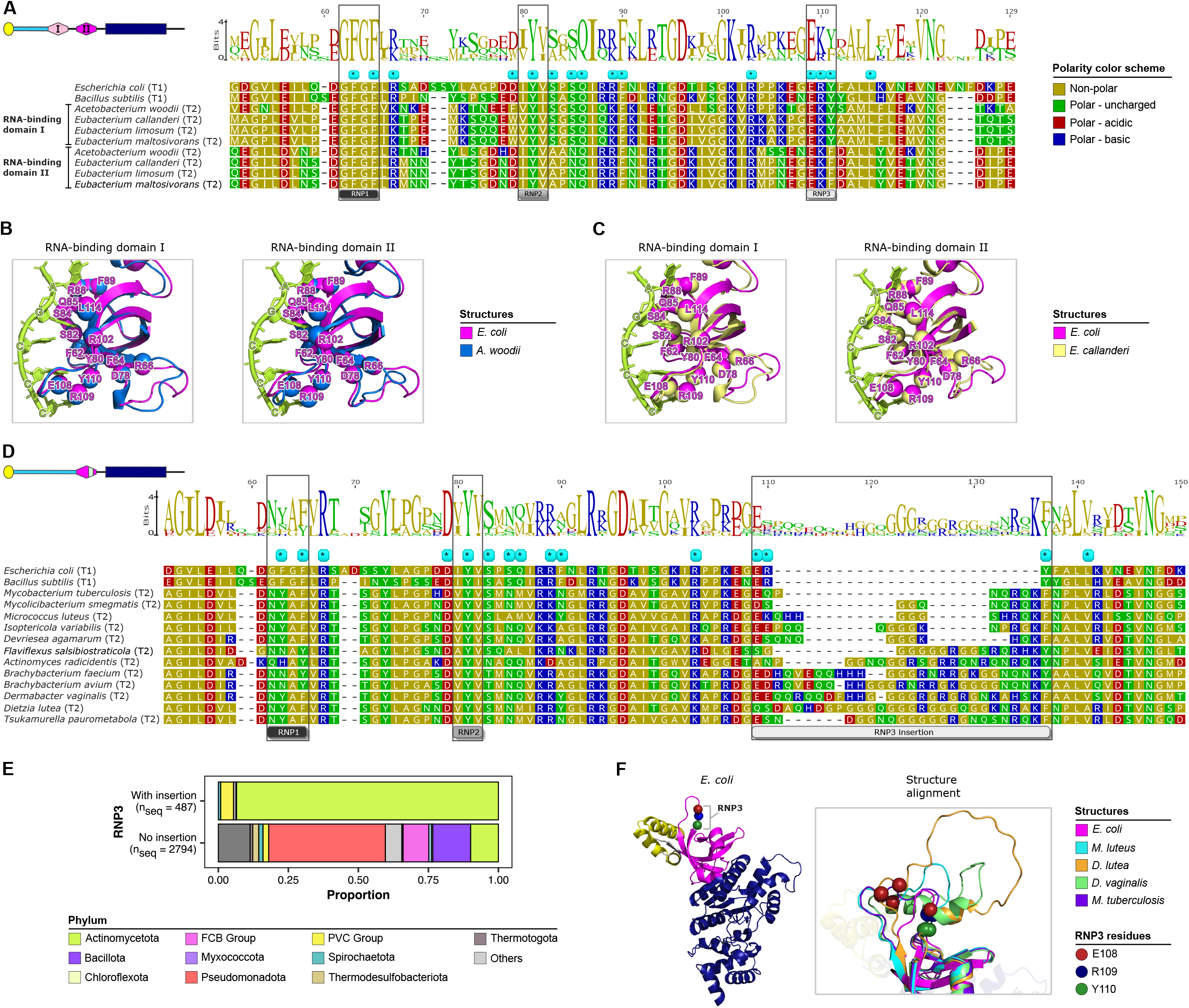
Variations in the Rho RNA-binding domain compared to _Ec_Rho. (A) Four species of Rho Type 2 from the Eubacteriaceae family (Bacillota) with duplicated adjacent RNA-binding domains. (B) Structure alignment of the RNA-binding domain of _Ec_Rho (PDB ID: 8E6W) with this duplicated domain in the AlphaFold structure of Rho from *Acetobacterium woodii* (AlphaFold ID: AF-H6LEI7-F1). (C) Structure alignment of the RNA-binding domain of _Ec_Rho (PDB ID: 8E6W) with this duplicated domain in the AlphaFold structure of Rho from *Eubacterium callanderi* (AlphaFold ID: AF-E3GFW7-F1). (D) Some Actinomycetota species with a small insertion in the RNA-binding domain in Rho Type 2 (T2). (E) Distribution of the RNP3 insertion among the bacterial phyla. (F) Secondary structure of _Ec_Rho (PDB ID: 8E6W) highlighting the RNP3 residues and the alignment of this structure with some Rho factors that have the RNP3 insertion. The sequence of _Ec_Rho and _Bs_Rho (*Bacillus subtilis*) were used as reference for the sequence alignments. Blue stars indicate the positions of the primary binding site in _Ec_Rho (PDB IDs: 1PVO and 8E6W). Alignments were performed by MUSCLE and colored according to the polarity scheme in Geneious Prime.

Another distinct feature in the RNA-binding domain was detected in some sequences of Type 2 Rho: an insertion of up to 20 amino acids containing enriched residues of arginine, glutamine, glycine, lysine, and glutamate (Table S8) with a conserved lysine residue at the C-terminal (Figure 5D and Figure S14). This small insertion was present mainly in Actinomycetota and the PVC Group (Figure 5E) and had previously been observed in *M. luteus* Rho (28) where it contains six basic residues in 14 amino acids (Figure 5D). Interestingly, those species do not have a conserved RNP1 (amino acids GFGF, Figure 5D), suggesting that the RNA-binding process might occur differently in these species. The structure alignment of some Rho AlphaFold predictions and _Ec_Rho (PDB ID: 8E6W) revealed that the RNP3 insertion could form a basic loop facing the exterior (Figure 5F). This position suggests the basic residues in the RNP3 insertion have a role in RNA binding. Notably, the genomes of these species have very high-GC genomes (62.2 % to 72.8 %) with RNA secondary structures that might require melting by additional RNA-binding regions (9,28,85).

### The Rho ATPase domain

The key regions of the ATPase domain of _Ec_Rho are Walker A and B motifs, Q- and R-loops, and the catalytic glutamate, arginine valve and arginine finger residues (88) (Figure 1A). Our new model is specific to Rho and incorporates BCM binding residues; however, it also detects the domain in species resistant to BCM. Overall, our analysis showed that this domain is highly conserved among the functional Rho factors (Figure S15). Short insertions (from five to 18 amino acids) in this domain were detected in 30 sequences mainly from the Acetobacteraceae family (Pseudomonadota) and a few members of Chloroflexota and Planctomycetota (Figure S16). These extra residues do not alter any essential Rho ATPase motif. Deletions (from five to 28 amino acids) were found in 203 sequences from several phyla, located at the very beginning or the end of the Rho ATPase domain (Figure S17). This observation could be due either to biases of our developed Rho ATPase model in identifying the exact position of this domain, or to loss of non-essential residues in this domain.

The conservation level and the fact that large insertions or deletions were not detected in this domain confirm the importance of secondary RNA-binding, helicase, and translocase activities of the ATPase domain when Rho terminates transcription. AlphaFold predictions suggest that smaller insertions are located between secondary structural elements, and they do not disrupt the overall ATPase domain fold (Figure S18). Consistent with this, point mutations at this domain in _Ec_Rho led to defective termination (16,19,94–96).

### BCM binding residues

BCM has activity against many species and was recently reported to act against clinically drug-resistant strains of *E. coli, Klebsiella pneumoniae,* and carbapenem-resistant Enterobacteriaceae (CRE) (97). As resistance is associated with mutations in the BCM binding pocket (83,98–101), species bearing similar substitutions in the BCM-binding pocket may be resistant to BCM.

We searched for variation in residues in the established BCM-binding interacting residues in _Ec_Rho (E211, R212, D265, S266, R269, T323, and G337) (34) and P180, K184, D210, L320 predicted to interact by Arpeggio (67) in the updated structural model of *E. coli* Rho and BCM (PDB ID: 1XPO). Notably, L320M was previously the only change found at L320 and confers BCM-resistance in *M. tuberculosis* (83). Substitutions at the previously known residues were found in 294 sequences, mainly from Rho with additional regions (Rho Type 2) (Table S9). Substitutions known to confer BCM resistance in *E. coli* (S266A and G337S) were detected mainly in *Rickettsia*, and Rho Type 2 proteins from *Fusobacterium* and *Streptomyces*, as observed previously (23). We observed the L320M substitution in all Mycobacteria and in several species from the order Corynebacteriales, as described by Saridakis et al. (82). We also observed the substitutions in P180A, P180S, P180Q, K184R in five species of Actinomycetes. BCM-binding pocket mutations may be important for providing resistance in species that produce BCM, and there are few examples of BCM-producing species that also have BCM binding pocket mutations: *Streptomyces formicae*, *S. kanamyceticus*, *S. platensis* (G337S) and *Mycobacteroides chelonae* (L320M) (Table S9 in bold) (102). But since there are other resistance mechanisms (103,104), we would not expect a perfect correlation. In *Staphylococcus* species, there was a substitution of D210G; however, in *Staphylococcus aureus* Rho is not essential and Rho is inhibited by BCM (25). Therefore, D210G does not confer BCM resistance in this context.

Substitutions in the BCM binding pocket could hinder BCM binding but may also reduce Rho efficiency (83). To characterize variations that may affect BCM binding we utilized the 749 predicted AlphaFold structures that were available for Rho. As these predictions do not include ligands, including BCM, we inserted of BCM into each Rho model using Alphafill (66) based on structure of the *E. coli* complex. The known L320M change for *M. canetti* shown in Figure 6A would disrupt H-bonding and cause steric hinderance. Novel substitutions in other species from L to R or T in *Nocardioides* and V in *Pimelobacter* could have similar effects (Figure 6B). In addition, changes in P180 (to A, S, Q) might affect the conformation of the binding pocket (Figure 6C). These data highlight species where BCM may be ineffective.

**Figure 6:**
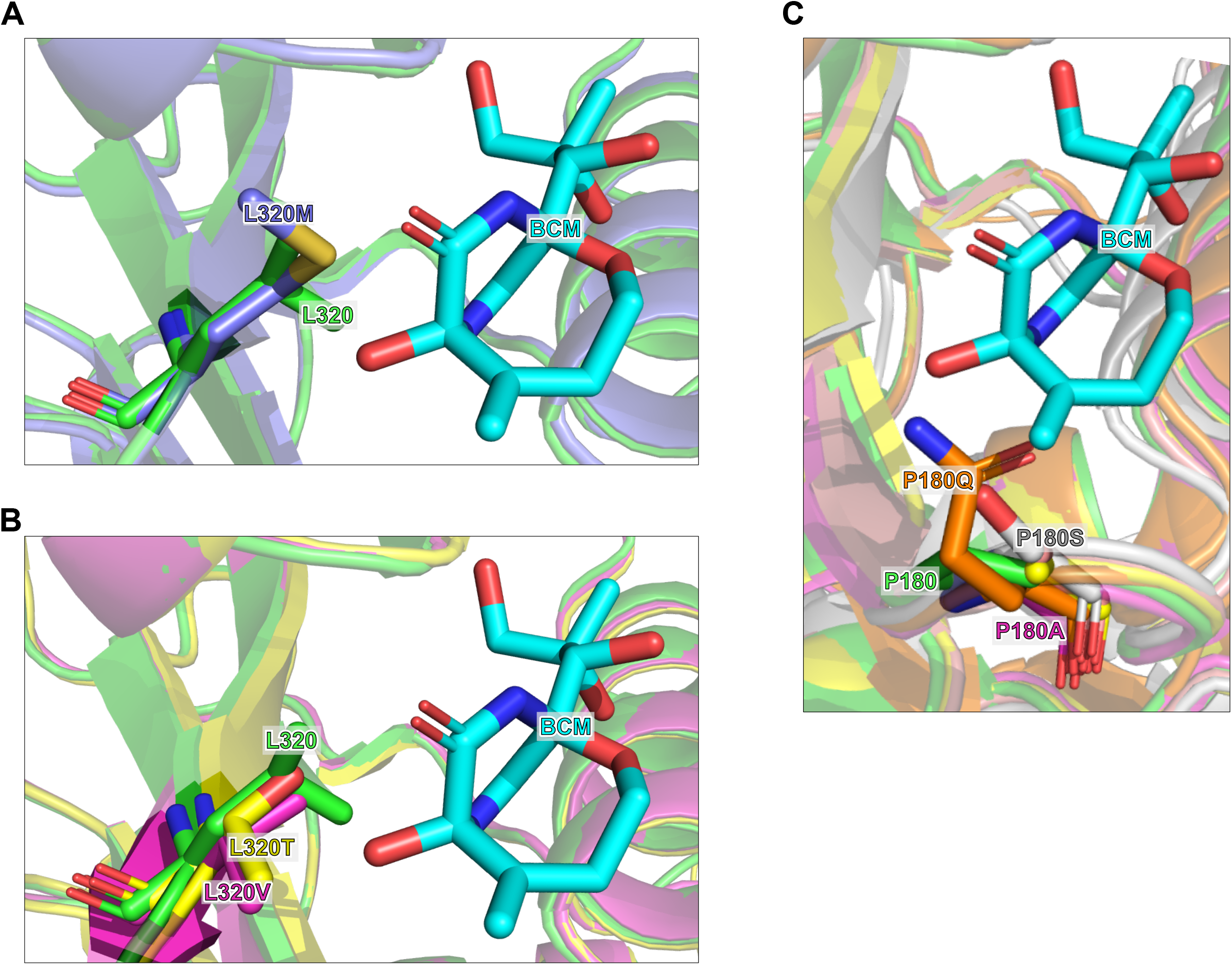
Substitutions in the BCM binding pocket could hinder BCM binding. (A) Known L320M in *M. canettii* (violet, AlphaFold ID: AF-A0A0K2HUQ9-F1). (B) Newly described L320T in *Nocardioides dokdonensis* (yellow, AlphaFold ID: AF-A0A1A9GPY2-F1) and L320V in *Pimelobacter simplex* (pink, AlphaFold ID: AF-A0A0A1DNV6-F1). (C) Newly described P180A in *P. simplex* (pink), P180Q in *Aeromicrobium erythreum* (orange, AlphaFold ID: AF-A0A0U4CG35-F1), and P180S in *Tropheryma whipplei* (grey, AlphaFold ID: AF-Q83MV6-F1). The Rho 3D structures were retrieved from AlphaFold and the BCM structure was retrieved from PDB (PDB ID: 1XPO). All panels have _Ec_Rho in green. Residues were colored in PyMOL.

### The three additional regions in Rho have diverse sequences suggestive of function

After studying the sequence of the three main domains, we analyzed the sequences of the additional regions of Rho present in our dataset. The initial, insertion, 2d, and extension regions were detected in 18.7%, 26.6%, 7.8%, and 1.3% of the Rho sequences, with lengths up to 352, 455, 393, and 69 amino acids, respectively. The initial and insertion regions are expected to be positioned near the primary RNA binding sites on the outside of the hexamer (Figure 1B, graphical abstract). Actinomycetota was the phylum with the highest proportions of sequences containing the initial (47.7%) and/or insertion (80.3%) while the extension (93%) was found mainly in Bacillota but only in 9.4% of the sequences. About 69% of the FCB Group had an insertion region. In the PVC Group and Spirochaetota, nearly half of the sequences had the initial or the 2d regions.

The additional regions have variable lengths and repeated compositional biases (e.g., high Q, R, or N content) (Figure 7, Figure S19, and Figure S20). Standard alignments of the full regions with ClustalW (105) or MAFFT (42) did not provide useful information about the conservation of these sequences or the presence of distinct motifs between more distantly related species (Figure S19). Given the overall dissimilarity and distribution it is likely these have arisen separately in different lineages (Figure 3). However, they may have common regions or functions.

**Figure 7:**
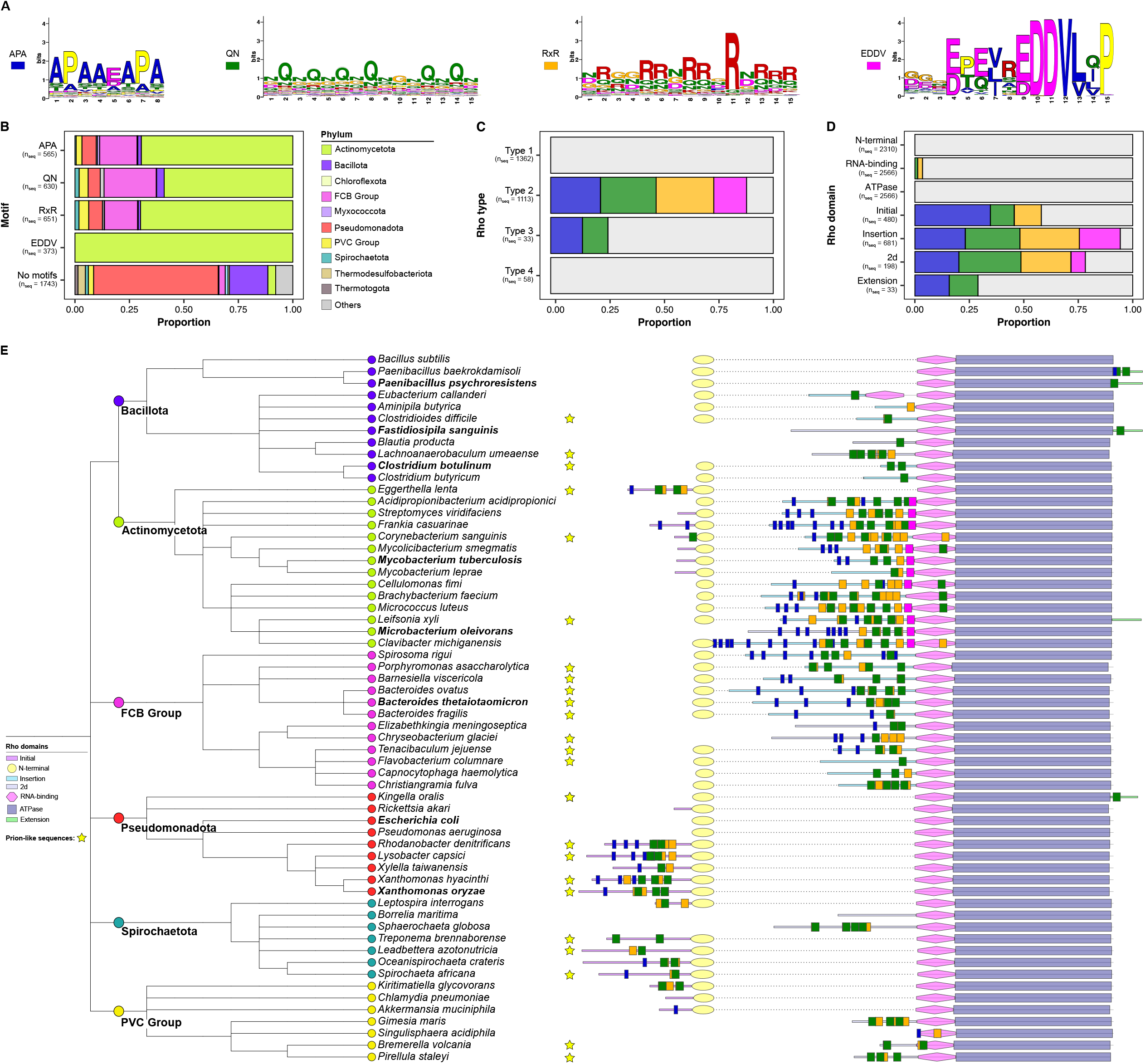
Rho additional regions have common motifs. (A) Sequence logos of the significant motifs. (B) Distribution of the significant motifs among the bacterial phyla. (C) Distribution of the significant motifs among the Rho types. (D) Distribution of the significant motifs among the Rho domains. (E) Phylogenetic tree based on the NCBI taxonomy of 60 species showing the distribution of the significant motifs detected in the Rho additional regions. Motifs were predicted by MEME and identified with MAST. Sequence logos represent the significant motifs. Prion-like sequences are highlighted with yellow stars. Species in bold have the Rho sequence characterized in detail in Figure 8.

The compositional bias of the insertion region, which was called IDR (intrinsically disordered region), of Rho in a large number of sequences was analyzed by Schumbera et al. (106). Arginine, aspartic acid, asparagine, and glycine were found enriched in the insertion region, but no common regions were defined (106). Hence, we employed the MEME Suite Tools to delineate smaller enriched motifs within the additional regions. Four significantly enriched motifs were identified (Figure 7A): (i) an “APA” motif: widespread in the initial and extension regions, particularly at the beginning of the insertion region; (ii) a “QN” motif: broadly detected in the center of the initial and insertion regions; (iii) a “RxR” motif: mostly found in the center and the C-terminus of the insertion region; (iv) an “EDDV” motif: only discovered at the end of the insertion region (adjacent to the RNA-binding domain) in Actinomycetota (Figure 7B).

Among the types, the four motifs were only detected in Rho Type 2. Rho Type 3 have some sequences with the APA and QN motifs while the Rho Type 1 and Type 4 do not show any occurrence (Figure 7C). Similarly, none of the four motifs are found in the N-terminal or the Rho ATPase domains. The few matches of the QN and the RxR motifs in the RNA-binding domain are in the RNP3 insertion (Figure 7D). Moreover, the RNA-binding domain with alterations (RNP3 insertion) is associated with additional regions with the RxR motif (Table S10).

When these presence and order of these motifs were examined within a phylogenetic context in the Rho additional regions, certain patterns emerged (Figure 7E and Figure S20). The Rho additional regions within Actinomycetota display a distinctive composition with the pattern of APA, QN, RxR, and EDDV motifs, with the EDDV only in the insertion region, as mentioned above. Those from Bacillota, Pseudomonadota, and the FCB Group also have APA, QN, RxR motifs (Figure 7B). All the extension sequences, even from different phyla (Bacillota, Actinomycetota, and Pseudomonadota), are grouped together showing the APA and QN motifs (Figure 7E). We also note a group of initial sequences from the Xanthomonadales order (Pseudomonadota) that have repeated copies of the APA and QN motifs (Figure S20).

The unusual composition and length variability of Rho additional regions suggest that those regions offer extra functions and properties to the typical Rho factor. For example, the motif APA could confer increased hydrophilicity and hydrodynamic volume, as found in the novel class of biosynthetic polymers called proline/alanine-rich sequence (PAS) polypeptides (107). Furthermore, the presence of alanine and proline-rich content is linked with disordered proteins (108), implying that the additional regions harboring multiple copies of these residues lack distinct secondary structures. In such cases, atypical Rho factors could undergo conformational changes exposing or hiding motifs/domains, potentially contributing to bacterial adaptability under environmental stresses.

The presence of the QN motif in Rho additional regions may also promote alternative structures such as in yeast prion-forming proteins. Most prion-forming proteins assemble as amyloid aggregates due to prion-forming domains that are intrinsically disordered and enriched in asparagine (N) and/or glutamine (Q) residues (109,110). In fact, the atypical Rho from *C. botulinum* has a highly polar, asparagine-rich insertion region capable of spontaneously self-assembling into amyloid-like structures *in vitro* (29) (Figure 7E). The aggregated form of *C. botulinum* Rho in *E. coli* showed compromised termination and genome-wide readthrough of Rho-dependent terminators (32). However, the presence of the QN motif is not sufficient to make a protein sequence a prion (111). In our study, only some additional regions with that motif were also predicted to be prion-like (yellow stars in Figure 7E and Figure S20). Rho from *B. fragilis* also has an insertion region with the QN motif (bold in Figure 7E), but this protein was not found to form aggregates in the conditions that were tested (30).

Besides conformational changes, the Rho additional regions may offer extra sites for RNA-binding. Arginine-rich motifs interact with RNA with notable affinity and specificity (112). Similarly, the arginine-rich motif (RxR) detected in the initial and insertion regions could bind to RNA, acting as extensions of the N-terminal and/or RNA-binding domains to assist in the recruitment of *rut* sites. Indeed, Rho from *M. tuberculosis* and *M. luteus* have arginine-rich insertion domains that were experimentally found to increase Rho affinity for RNA and to help promoter-proximal RDT, similar to NusG (28,82). Another possible extension of the RNA-binding domain is the unique EDDV motif found immediately before it (Figure 7E, Figure S20). Although there are no similar motifs reported in the literature, this motif is likely to have an important function given its high degree of conservation in the insertion regions of Actinomycetota, and its location adjacent to the RNA-binding domain.

Arginine-rich motifs are also associated with intrinsically disordered proteins and phase separation (113,114). Therefore, those features could be present in the atypical Rho factors with the RxR motif as demonstrated in Rho of *B. thetaiotaomicron* (26). Schumbera et al. (106) reported amino acid bias in the insertion region; we find a richer set of more complex motifs in insertion also in the initial and extension regions (Figure 7).

### The additional regions are predicted to be intrinsically disordered and to undergo phase separation, with some having prion-like portions

The insertion regions of Rho from *C. botulinum* and *B. thetaiotaomicron* have been experimentally shown to exhibit prion-like characteristics and induce phase separation, respectively (26,29,32). The type of sequence motifs observed have been associated with prion-like characteristics and phase separation, we therefore predicted the likelihood of prion-like properties and/or intrinsic disorder in these regions. This was done for four experimentally tested Rho (Figure 8A to D) and those predictions were expanded to those from *Xanthomonas oryzae* (with initial region) (Figure 8E), *Paenibacillus psychroresistens* (with extension region) (Figure 8F), *Microbacterium oleivorans* (with 2d region) (Figure 8G).

**Figure 8:**
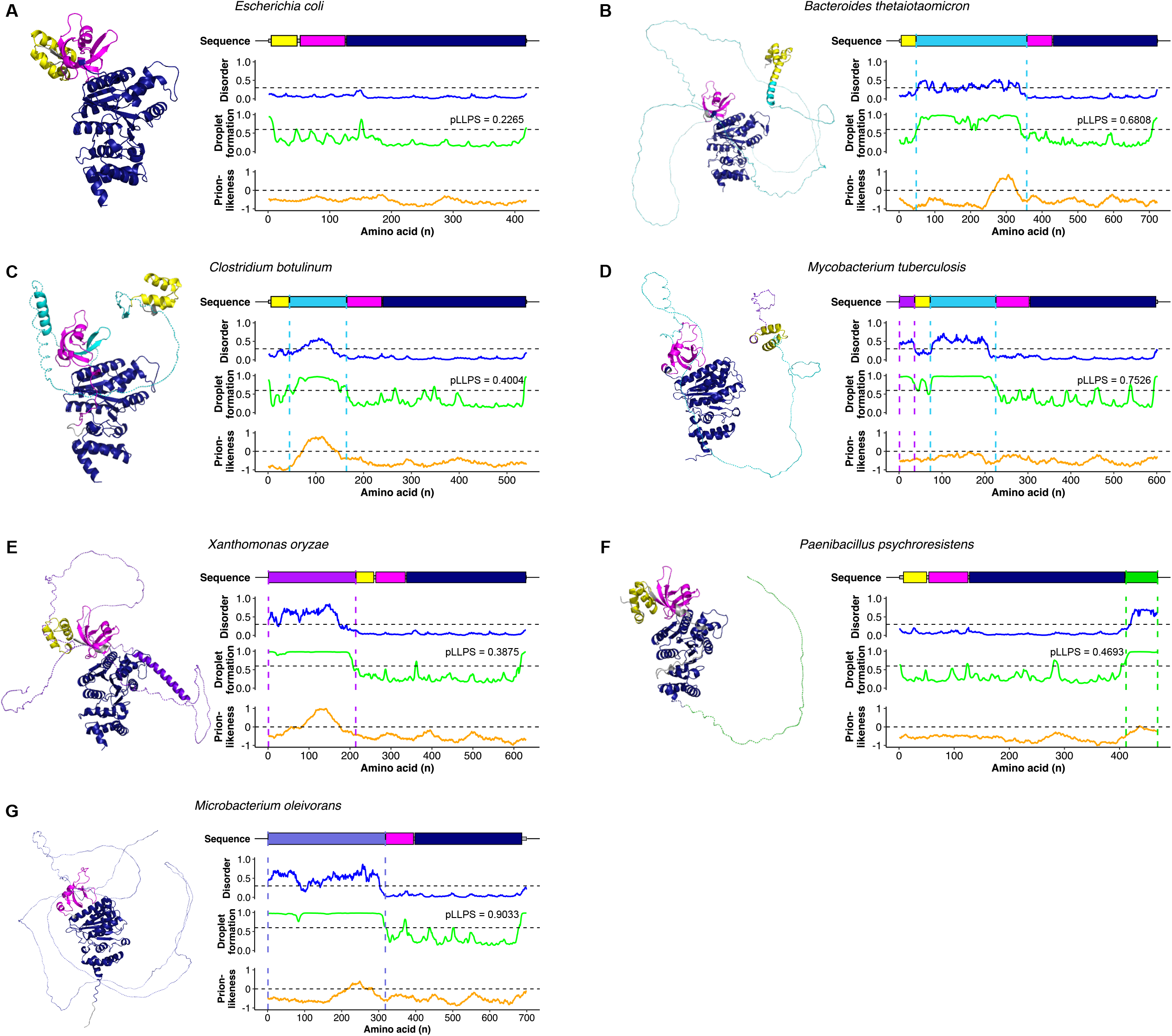
Characterization of selected bacterial Rho factors for disorder, droplet formation, and prion-likeness. (A) Typical Type 1 Rho from *E. coli* K12 (AlphaFold ID: AF-P0AG30-F1). (B) Atypical Type 2 Rho with an insertion region (+309 aa) from *Bacteroides thetaiotaomicron* DSM 2079 (AlphaFold ID: AF-C6ITN0-F1). The insertion regions in panels B - E are predicted as loops by AlphaFold with very low confidence pLDDT <50. Very low scores are indicative of disorder (123). (C) Atypical Type 2 Rho with an insertion region (+119 aa) from *Clostridium botulinum* E1 str. ’BoNT E Beluga’ (AlphaFold ID: AF-C4IEE6-F1). (D) Atypical Type 2 Rho with both initial (+35 aa) and insertion (+152 aa) regions *Mycobacterium tuberculosis* H37Rv (AlphaFold ID: AF-P9WHF3-F1). (E) Atypical Type 2 Rho with an initial region (+214 aa) from *Xanthomonas oryzae* pv. oryzicola (AlphaFold ID: AF-A0A0C5V4F0-F1). (F) Atypical Type 3 Rho with an extension region (+107 aa) from *Paenibacillus psychroresistens* (AlphaFold ID: AF-A0A6B8RTJ4-F1). (G) Atypical Type 2 Rho with the 2d region (+318 aa) from *Microbacterium oleivorans* (AlphaFold ID: AF-A0A031FP33-F1). 3D structures were extracted from the AlphaFold database and were colored according to the different domains. The additional regions were not confidently modelled (purple, light blue, and green dashed lines) possibly due to their disordered nature. Predictions were performed by flDPnn, FuzDrop, and PLAAC for disorder, droplet formation, and prion-likeness, respectively. Abbreviation: pLLPS, Probability of forming a droplet state through liquid-liquid phase separation. Proteins with pLLPS ≥ 0.60 are droplet-drivers (can spontaneously undergo liquid-liquid phase separation) and proteins with pLLPS < 0.60 but possess droplet-promoting regions are droplet-client proteins (can induce their partitioning into condensates).

The three main domains of the typical Rho factor from *E. coli* were not predicted to be disordered, promote droplet formation, or be prion-like (Figure 8A), consistent with the many available structures for _Ec_Rho in the PDB database (e.g., PDB IDs: 6WA8, IPV4, IPVO, 3ICE, and 8E3H) (5,6,34,88,115). In contrast, the insertion region of Rho from *B. thetaiotaomicron* was predicted to be intrinsically disordered and spontaneously undergo liquid-liquid phase separation, consistent with recent experimental observations (26) (Figure 8B). The disorder of the insertion region is consistent with the inability of Alphafold to predict the structure (Figure 8B). Additionally, the section of 62 residues with multiple copies of the QN motif (Figure 7), located at the C-terminus of the insertion region, was predicted to be prion-like (Figure 8B).

Similar to *B. thetaiotaomicron*, the insertion from *C. botulinum* was predicted to be disordered, undergo droplet formation, and be prion-like (Figure 8C). The overall probability of *C. botulinum* Rho to form a droplet state through liquid-liquid phase separation (pLLPS) was predicted to be 0.4, which falls below the threshold of 0.6 determined by FuzDrop (63). However, a short section of about 70 residues has a propensity to droplet formation and predicts this Rho factor is a droplet-client protein that could induce its partitioning into condensates. This short region also corresponds to the N rich self-assembly prion like region tested by Pallarès et al. (29) as well as Yuan and Hochschild (32).

The initial and insertion regions of the Rho from *M. tuberculosis* were not predicted to be prion-like (Figure 8D) but were predicted to be disordered and promote phase separation. The intrinsic disorder trait of the additional regions of Rho is consistent with the fact that Saridakis et al. (83) were not able to resolve the structure of the first 220 residues of each monomer of *M. tuberculosis* Rho; this region corresponds exactly to the initial, N-terminal and insertion regions.

Disorder and droplet formation were also predicted for the initial and the extension regions of Rho from *X. oryzae* (Figure 8E) and *P. psychroresistens* (Figure 8F), respectively. Moreover, the initial region in *X. oryzae* Rho had a 95-residue QN-rich sequence that was predicted to be prion-like. The 2d region in *M. oleivorans* (Figure 8G) had propensity to disorder and droplet formation.

Following the confirmation that the bioinformatics predictions align with the experimental data for Rho, we extended our analysis to encompass a representative range of non-redundant Rho sequences (n = 519). As prion-likeness was detected primarily in short sections within the additional regions, we conducted an analysis of the mean scores within 25-residue windows for each Rho domain to assess disorder (Figure 9A), droplet formation (Figure 9B), and prion-likeness (Figure 9C). The results showed that the Rho main domains (N-terminal, RNA-binding, and ATPase) did not have these propensities. In contrast, the initial, insertion, 2d, and some parts of the extension regions were predicted to be disordered (Figure 9A) and promote droplet formation (Figure 9B). Most of the Rho residues in all domains/regions do not display a propensity for prion-like behavior (Figure 9C). However, 16.7%, 10.4%, 14.0%, and 18.8% of windows in the four types of additional regions (initial, insertion, 2d, and extension, respectively) were predicted to display prion-like characteristics (PLAAC score >= 0). In contrast to the primary Rho domains, that exhibited 0 (N-terminal and ATPase) or 0.5% (RNA-binding) of windows with propensity to prion-like (Figure 9C).

**Figure 9:**
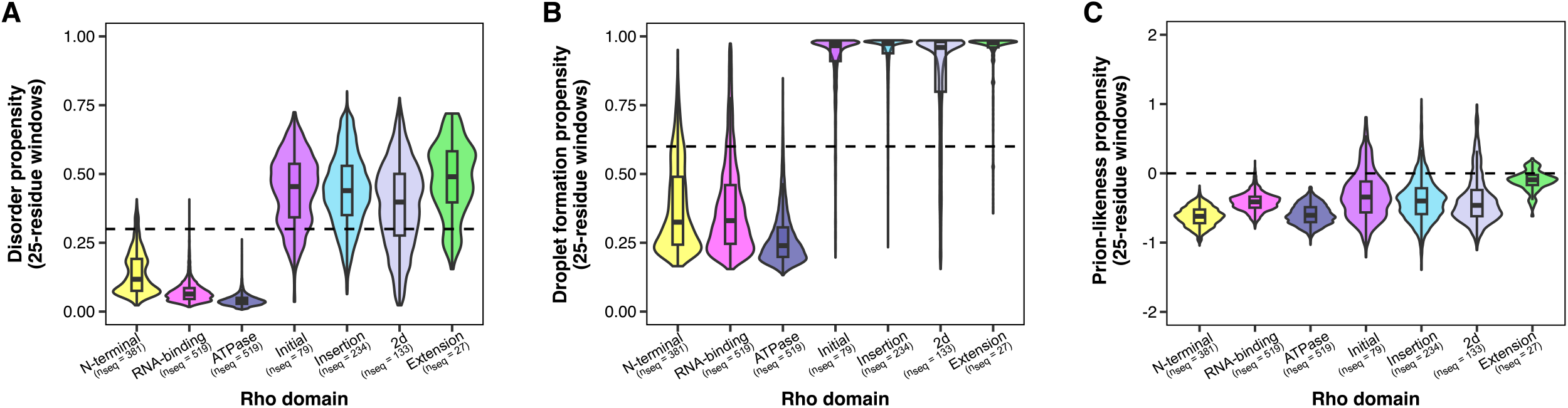
The additional regions of atypical Rho factors have predictions for disorder, droplet formation and prion-likeness. Violin and box plots showing the mean propensity of each Rho domain to: (A) disorder, (B) droplet formation, and (C) prion-likeness in 519 non-redundant sequences. Predictions were performed by flDPnn, FuzDrop, and PLAAC for disorder, phase separation, and prion-likeness, respectively. The classification threshold for each program is represented by the dashed line (flDPnn: 0.3; FuzDrop: 0.6; PLAAC: 0). Box plots depict the distribution of the mean of the predicted scores as quartiles. The line at the center of each boxplot corresponds to the median value. Violin plots represent the density distribution of the mean scores. Pairwise p-values are shown in Table S11.

Our predictions indicate that the properties of intrinsic disorder, phase separation, and prion-likeness are likely associated with the additional regions but not the main Rho domains. Based on the known interaction features of IDRs (116), we speculate that the extra properties confer additional properties on Rho. Since phase separation can be induced during stress conditions due to changes in the protein solubility (117), the formation of biomolecular condensates may concentrate atypical Rho factors, facilitating the binding of weak terminators. Indeed, liquid-liquid phase separation LLPS of the atypical Rho from *B. thetaiotaomicron* was recently observed upon starvation (26), and prion behavior was perceived in the atypical Rho of *C. botulinum* upon ethanol stress (32). Moreover, evidence of LLPS of proteins contributing to bacterial fitness has been shown for *Caulobacter crescentus* (118,119) and *E. coli* (120–122).

## CONCLUSIONS

Upon analyzing a larger set of high-quality bacterial genomes (n = 2,730), using specific models, and applying recently developed bioinformatics tools, we greatly extended previous functional predictions for parts of Rho.

In addition to the typical Rho factor observed in *E. coli* and the insertion region described in previous studies across different bacteria, our study has uncovered a more complex landscape. Specifically, we have detected initial and extension regions within the Rho factor, leading to the identification of four distinct major architectural types of functional Rho.

The architectural diversity of Rho is not only a structural discovery but potentially indicates that the additional regions are functionally equivalent. This inference is demonstrated by the identification of common motifs within these additional regions, providing the Rho factor with extra properties. These properties, which could be enhanced RNA-binding capabilities and the ability to undergo conformational changes due to attributes like intrinsic disorder, phase separation, and prion-like behavior, hold implications for Rho’s function, notably, the promotion of fitness during periods of environmental stresses.

## DATA AVAILABILITY

All bacterial genomes were taken from public databases. The data used are provided in the article and in the supplemental material. Any further requests can be sent to.

---

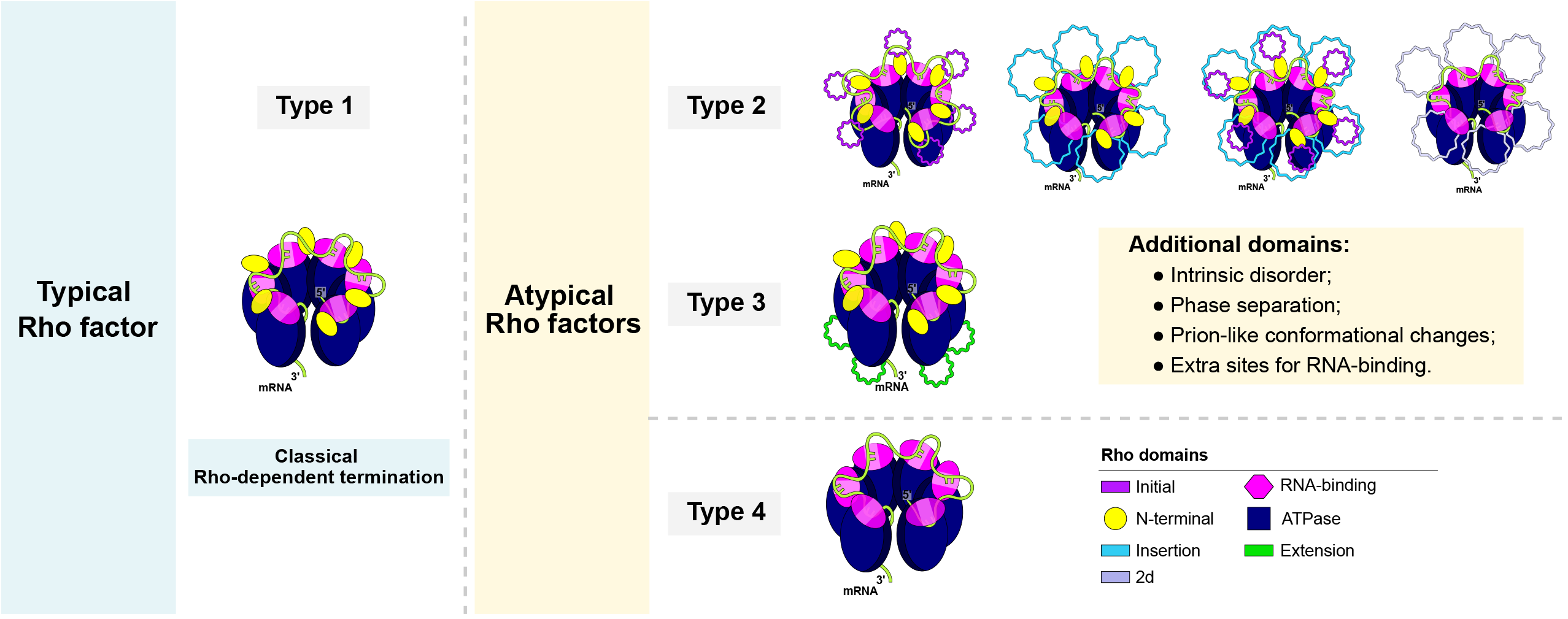

