## Supplementary Material for "Diversification of the Rho transcription termination factor in bacteria"

**Supplementary Text**

**Species lacking all Rho domains**

A total of 232 bacterial species did not have any Rho domain, and they comprise 90% (n = 90) of Mycoplasmatota, 29% (n = 141) of Bacillota, and one member of Cyanobacteriota (*Prochlorococcus marinus* subsp. *marinus* str. CCMP1375). From the Bacillota phylum, species completely devoid of Rho were: (i) Negativicutes such as *Acidaminococcus fermentans* and *Selenomonas sputigena*; (ii) Desulfitobacteriaceae such as *Desulfitobacterium dehalogenans* ATCC 51507 and *Desulfosporosinus acidiphilus;* (iii) Streptococcaceae including all *Streptococcus* with exception of *Streptococcus pasteurianus* and *Streptococcus pseudopneumoniae* IS7493 that contained Rho-like N-terminal domain sequences; (iv) most members of Lactobacillaceae excluding *Pediococcus* sp., *Lactiplantibacillus* sp., *Ligilactobacillus* sp., and *Loigolactobacillus* sp.

**Species containing Rho domains**

There are four major structural types of functional Rho factors, which we named as Type 1 (typical Rho with the three main domains), Type 2 (atypical Rho with an additional initial and/or insertion region), Type 3 (atypical Rho with an additional extension region), and Type 4 (atypical Rho lacking the N-terminal domain but containing the RNA-binding and the Rho ATPase domains). Rho Type 2 can be further subdivided into four subtypes: Subtype 2a (atypical Rho with an initial region), Subtype 2b (atypical Rho with an insertion region), Subtype 2c (atypical Rho with both initial and insertion regions), Subtype 2d (atypical Rho with an initial/insertion - “2d” region).

In contrast, there are four possibilities of Rho-like sequences in which one or two of the main Rho domains are missing: Rho-like N-terminal domain, Rho-like RNA-binding domain, Rho-like N-terminal and RNA-binding domains, and Rho-like ATPase domain.

**Rho Type 1 (**
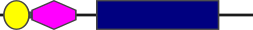
**)**

The typical Rho was the most abundant group with 1,363 sequences (44.7%) in 1,359 organisms (49.8%). It was mainly found in Pseudomonadota (n = 967), but not in Actinomycetota. Species containing duplicated sequences were: *Fervidobacterium pennivorans* DSM 9078 (Thermotogota), *Ktedonosporobacter rubrisoli* (Chloroflexota)*, Luteitalea pratensis* (Acidobacteriota), and *Terriglobus roseus* DSM 18391 (Acidobacteriota). The largest sequence (453 aa) was from *Ferrovibrio terrae* (Pseudomonadota). The smallest sequence (411 aa) was from *Thermaerobacter marianensis* DSM 12885 (Bacillota). Overall, Type 1 sequences were very conserved, and the three main domains had all expected essential residues.

**Rho Subtype 2a (**
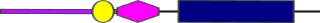
**)**

The atypical Rho with an initial region was found in 234 (7.7%) sequences of 226 (8.3%) organisms. This group was mainly detected in Pseudomonadota (n = 83), the PVC Group (n = 38) and Spirochaeota (n = 21), but not in the FCB Group. Only one member of Bacillota (*Limnochorda pilosa*) had Subtype 2a. Species containing duplicated sequences were the tick-borne pathogens of Rickettsiales order *Anaplasma* sp. and *Ehrlichia*. The largest sequence (711 aa) was from *Coriobacterium glomerans* PW2 (Actinomycetota) that had a long initial region with 352 residues. The smallest sequence (433 aa) was from *Pseudoleptotrichia goodfellowii* (Fusobacteriota) that had the initial region with 33 residues. Similar to Type 1, the three main Rho domains were well conserved. The initial region had multiple copies of the “APA” and “QN” motifs.

**Rho Subtype 2b (**
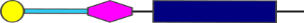
**)**

The atypical Rho with an insertion region was found in 435 (14.3%) sequences of 435 (15.9%) organisms. It was mostly detected in the FCB Group (n = 162), Actinomycetota (n = 175), and Bacillota (n = 91). Only one Pseudomonadota had Subtype 2b: the amylase-producing bacterium *Glaciecola amylolytica*. The largest sequence (849 aa) was from *Marisediminicola antarctica* (Actinomycetota) with an insertion of 422 residues. The smallest sequence (432 aa) was from *Hippea maritima* (Bacillota) that had the insertion region with 25 residues. The N-terminal and the Rho ATPase domains were well conserved. Ten species of Actinomycetota ([Actinomycetia](https://www.ncbi.nlm.nih.gov/Taxonomy/Browser/wwwtax.cgi?mode=Info&id=1760&lvl=3&lin=f&keep=1&srchmode=1&unlock" \o "class)) had an insertion H/Q/R rich of about 20 residues inside the RNA-binding domain. Four members of the Eubacteriaceae family in Bacillota (*Acetobacterium woodii* DSM 1030, *Eubacterium callanderi*, *Eubacterium limosum*, and *Eubacterium maltosivorans*) had duplicated adjacent RNA-binding domains. The first RNA-binding domain had more substitutions than the second one. However, both domains had the conserved primary RNA-binding residues. The insertion had multiple copies of the four significant motifs, and it was more conserved than the initial region.

**Rho Subtype 2c (**
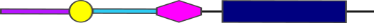
**)**

The atypical Rho with the two extra regions was found only in Actinomycetota (9% of the sequences, n = 246) without duplications. The largest sequence (972 aa) was from *Cryobacterium soli* with an initial region of 98 residues and an insertion region of 454 residues. The smallest sequence (585 aa) was from *Mycobacterium branderi* with an initial region of 42 residues and an insertion region of 454 residues. Overall, the N-terminal and the Rho ATPase domains were well conserved. Three species (*Brachybacterium vulturis, Tsukamurella paurometabola,* and *Tsukamurella tyrosinosolvens*) had a G/N-rich insertion inside the RNA-binding domain. The initial region was particularly enriched in the “APA” motif while the insertion region had all the four significant motifs following the pattern of “APA-QN-RxR-EDDV”.

**Rho Subtype 2d (**
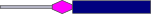
**)**

The pattern with the N-terminal domain not detected and having the RNA-binding domain placed after the 25^th^ residue was found in 198 (6.5%) sequences in 189 (6.9%) of the bacterial species. It was widely distributed in the main bacterial phyla: Actinomycetota (n = 53), FCB Group (n = 56), Bacillota (n = 15), PVC Group (n = 38), and Spirochaetota (n = 20). The longest sequence (776 aa) was from *Microbacterium lemovicicum* (Actinomycetota) with a 2d region of 394 residues. The smallest sequence (382 aa) was from *Actinoplanes missouriensis* (Actinomycetota) with the 2d region of 33 residues. Duplicated sequences with different lengths were found only in 12 species of the PVC Group such as *Akkermansia glycaniphil* and *Mariniblastus fucicola.* The RNA-binding domain was quite conserved excepting two Actinomycetota (*Actinotignum schaalii* and *Phytohabitans flavus*) that had an N/R rich insertion. The Rho ATPase domain was very conserved. Similar to other initial and insertion regions, multiple copies of the significant motifs were found in the 2d region including the “EDDV” motif in Actinomycetota species. Therefore, the 2d region may have the same functional properties as the other Rho additional regions.

**Rho Subtype 3 (**
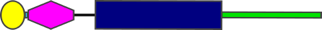
**)**

The atypical Rho with an extension region was detected in 33 (1.1%) sequences of 33 (1.2%) organisms. It was predominantly found in Bacillota (n = 32) (*Paenibacillus* sp., *Cohnella abietis,* and *Desulforamulus*) and in one species of Pseudomonadota (*Kingella oralis*). The largest sequence (482 aa) was from *Paenibacillus baekrokdamisoli* (Bacillota) with an extension of 69 residues. The smallest sequence (432 aa) was from *Cohnella candidum* (Bacillota) with an extension of 26 residues. Altogether, all the three main domains were well conserved. The extension region was conserved containing only the ”APA” and “QN” motifs.

**Rho Subtype 4 (**
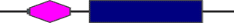
**)**

The group lacking only the N-terminal and with the RNA-binding domain located within the first 25 residues of the sequence was present in as a single copy of 58 (2.1%) organisms. It was mostly detected in Actinomycetota (n = 46) and a few members of the PVC Group (n = 5). The longest sequence (406 aa) was from *Catenulispora acidiphila* (Actinomycetota). The smallest sequence (341 aa) was from *Lentzea guizhouensis* (Actinomycetota). The RNA-binding domain was slightly divergent excepting the RNP residues. The Rho ATPase domain was quite conserved with the essential residues of _Ec_Rho. Given the essentiality of the RNA-binding and the Rho ATPase domains, those sequences could possibly terminate transcription conferring the classification of atypical Rho factors, rather than Rho-like proteins.

**Species with more than one copy of Rho: Group A and B Rho**

A previous study had noted that diverse species contained more than one copy of the Rho gene and that the second copy was generally shorter (named A and B) (1). Some of these were duplicates of A or B. We built two hmm models to distinguish the A and B (Methods, conserved regions shown in Figure S8). Phylogenetic analyses using the common part of the Rho (RNA-binding and Rho ATPase) showed four clearly distinct clusters of the pairs of sequences (Figure S6, central ellipses). There are two clusters (#1 and #2) in Actinomycetota (Fig S6, light green) that have dissimilar A and B Rho’s corresponding to Type 2 (cluster #1, Group A) and Type 4 (cluster #2, Group B), respectively. The remaining phyla had each copy of Rho separated in two clusters of Type 2d or Type 4 (cluster #3, Group B) and Type 2 (cluster #4, Group A). When the pairs were compared species by species (Figure S7), the first copy (larger sequence) is a Group A/Type 2, and the second copy (shorter sequence) is a Group B Rho without the N-terminal domain (Type 2d or Type 4) (Figure S7). The few exceptions are the two species of Acidobacteriota (*Terriglobus roseus* and *Luteitalea pratensis*) that have two distinct Type 1 sequences and two genera of Pseudomonadota (*Anaplasma* sp. and *Ehrlichia* sp.) that have exact Rho Type 2 duplications.

These data are consistent with a model where the Rho gene has duplicated anciently in Actinomycetota and separately other phyla. As the N-terminal domain lacking (Group A, or Type 4 Rho) are often present with a Rho with additional domains, it is possible that the additional domains provide the initial non-specific RNA binding function in trans in a heterohexamer.

**Rho-like N-terminal domain (**
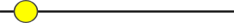
**)**

The Rho N-terminal domain is a bundle of three alpha helices containing in _Ec_Rho basic residues that bind RNA. This structural motif usually without the conserved amino acids of Rho is found alone and in other proteins. Sequences containing only the Rho N-terminal domain are named by NCBI as “Rho termination factor N-terminal domain-containing protein”, and it is not a functional Rho factor. It was present in 452 (14.8%) sequences in 395 (14.5%) organisms being found in all representative phyla excepting Spirochaetota. This group was mostly present in Actinomycetota (n = 198) and Pseudomonadota (n = 59). About 97% (n = 32) of Cyanobacteriota had Rho-like N-terminal domain sequences with 11 species having duplicated copies of those sequences. Three species had three copies of these sequences: *Mycolicibacterium aichiense* (Actinomycetota), *Oscillatoria acuminata* PCC 6304 (Cyanobacteriota), and *Gimesia maris* (PVC Group). The longest sequence (897 aa) was from *Limnospira fusiformis* SAG 85.79 (Cyanobacteriota), and the smallest sequence (33 aa) was from *Clostridium sporogenes* (Bacillota). Overall, these sequences were very divergent and might not be related to Rho.

**Rho-like RNA-binding domain (**
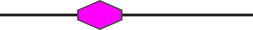
**)**

The sequence containing only the Rho RNA-binding domain is named by NCBI as “Rho RNA-BD domain-containing protein”, and it is not a functional Rho factor. It was present only in *Alistipes dispar* and *Alistipes megaguti* (Bacteroidota) with sizes of 435 and 434 aa, respectively. The RNA-binding domain was located at the first half of the protein (55-120). Some residues of the RNA-binding motifs are conserved.

**Rho-like N-terminal and RNA-binding domains (**
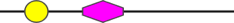
**)**

The sequences lacking the Rho ATPase domain are probably not functional Rho factors. This group was only found in two Actinomycetota of our dataset: *Baekduia soli* (Rubrobacteria Class) (472 aa) and *Conexibacter woesei* DSM 14684 (Thermoleophilia Class) (449 aa). Compared to _Ec_Rho, the N-terminal domain had the basic residues and also an insertion containing the “RxR” motif. The RNA-binding domain was quite conserved. Extending our analysis by investigating these sequences in related organisms from the Rubrobacteria and Thermoleophilia Classes, it is possible to observe that these Actinomycetota have valine or adenine instead of the catalytic glutamate and they lack the secondary binding sites (Q- and R-loops) compared to _Ec_Rho. Therefore, these Rho-like proteins would not be able to perform the traditional step of RNA secondary binding and RNA translocation to end transcription. The only exception was found for *Rubrobacter aplysinae* (Rubrobacteria Class) which has the Q- and R-loops.

**Rho-like ATPase domain (**
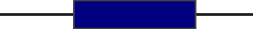
**)**

The sequences containing only the ATPase/BCM-binding domain are not functional Rho factors. This group was present in 28 (0.9%) sequences in 28 (1.0%) organisms. It was found only in Pseudomonadota such as *Colwellia psychrerythraea* 34H and *Marinomonas arctica,* the PVC Group (*Lacunisphaera limnophila*), and Chloroflexi (*Ktedonosporobacter rubrisoli*). The longest sequence (530 aa) was from *Corallococcus coralloides* DSM 2259 (Pseudomonadota) and its predicted secondary structure is similar to the typical Rho. The smallest sequence (282 aa) was from *Bradymonas sediminis* (Pseudomonadota). In general, sequences were well conserved and had the essential residues. Sequences from the Myxococcocales order have an initial domain D/G/R rich.

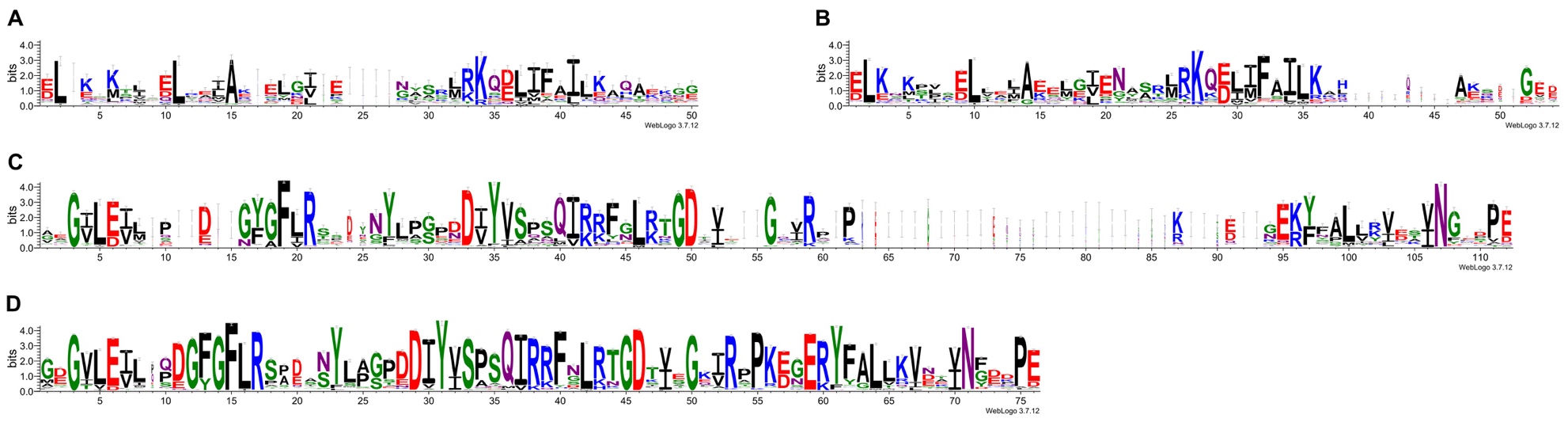

**Figure S1: Conserved features of the newly developed models for the Rho N-terminal and RNA-binding domains are similar to Pfam Rho models. (**A) Rho N-terminal domain model from Pfam (PF07498.16). (B) Developed model for the Rho N-terminal domain. (C) Rho RNA-binding domain model from Pfam (PF07497.16). (D) Developed model for the Rho RNA-binding domain. Sequence logos were generated from the sequence alignments on the WebLogo 3 server (https://weblogo.threeplusone.com).

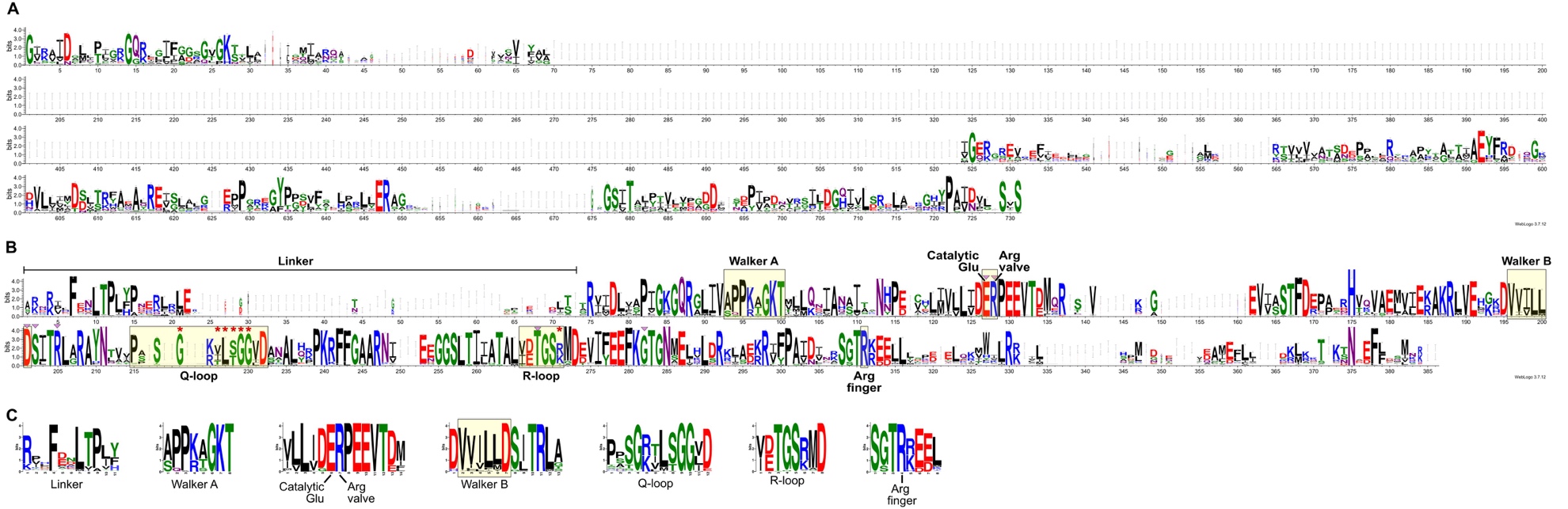

**Figure S2: The developed model for the Rho ATPase domain contains all characterized Rho motifs.** (A) Alignment of the sequences present in the Pfam ATPase model (PF00006.29). (B) Alignment of the sequences present in the Rho ATPase developed model. (C) MEME significant motifs detected on the sequences of the Rho ATPase developed model. Pink triangles and red stars highlight the residues forming the BCM-binding pocket and the secondary binding site in _Ec_Rho (PDB ID: 1XPO), respectively. Sequence logos were generated from the sequence alignments on the WebLogo 3 server (https://weblogo.threeplusone.com).

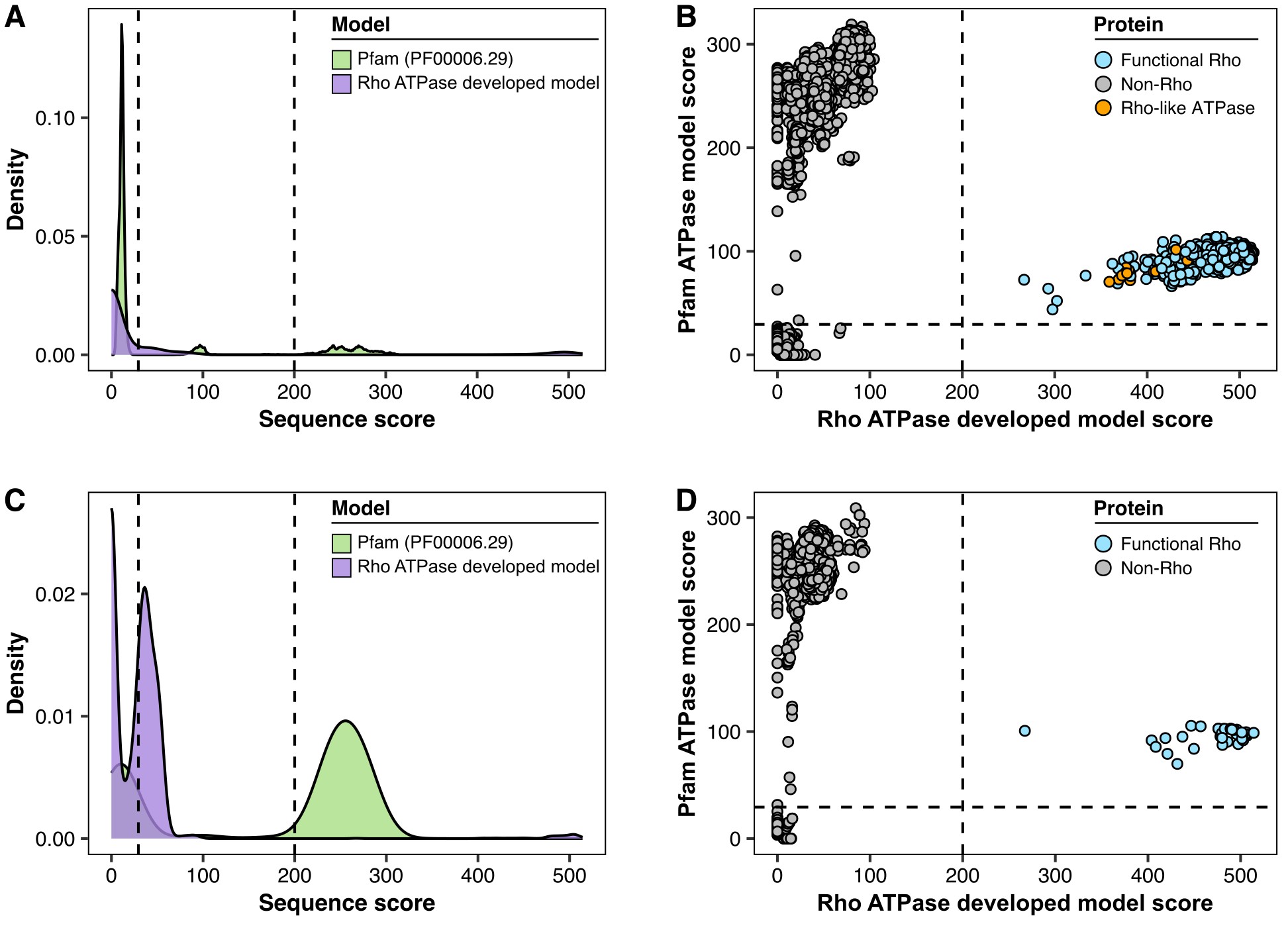

**Figure S3: Comparison between the ATPase models from Pfam (PF00006.29) and the Rho ATPase developed model.** (A) and (B): results for all bacterial protein sequences in our database. (C) and (D): results for the UniProtKB/Swiss-Prot protein sequence database. (A) and (C): density plots of the sequence scores for each model. (B) and (D): model scores plotted against each other. The dashed lines represent the threshold scores of the Pfam (29.4), and the Rho ATPase developed model (200). Sequence scores were calculated by hmmsearch.

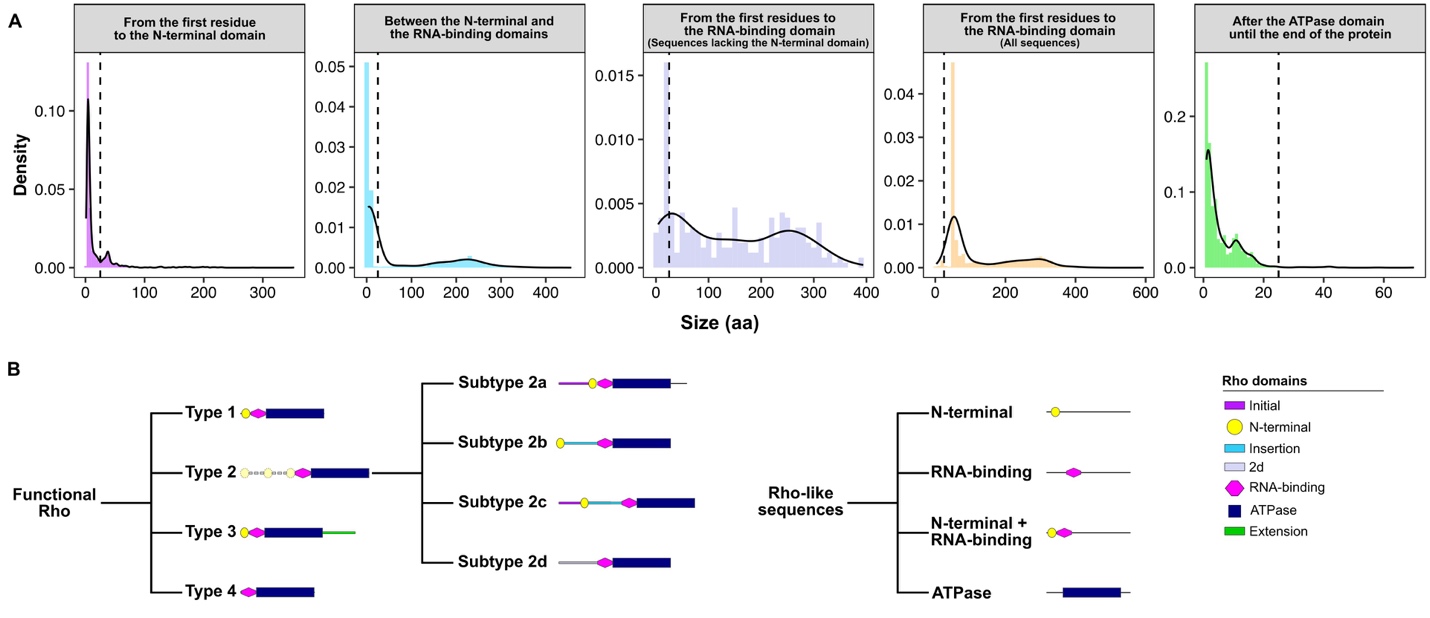

**Figure S4: Size of the Rho additional regions and representation of Rho sequence types.** (A) Density plots of the size of the regions between the three Rho main domains. The additional regions initial (purple), insertion (blue), 2d (light purple), and extension (green) are by definition at least 25 aa long (dashed line). (B) Graphic representation of the types and subtypes of the Rho as well as the Rho-like sequences.

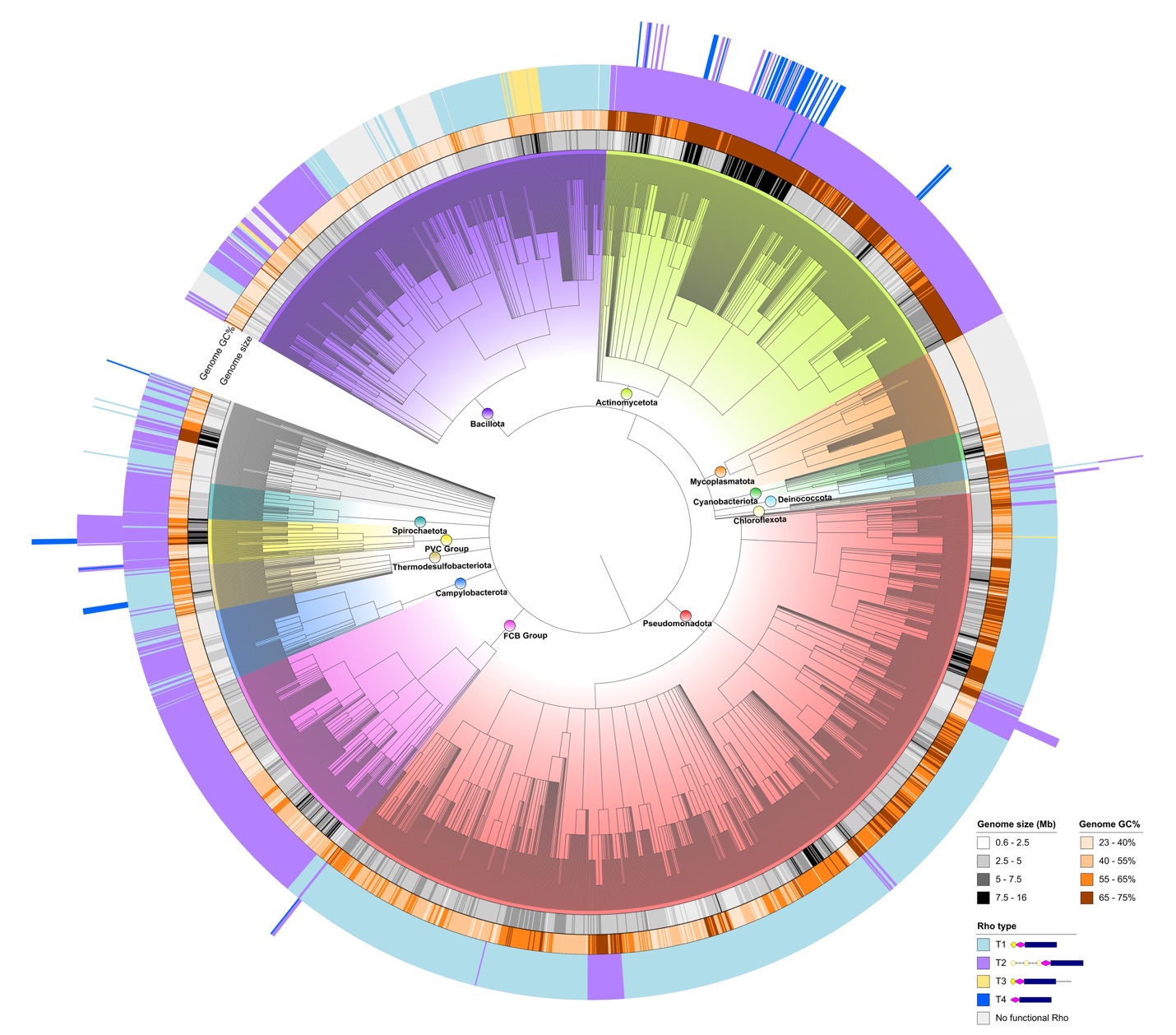

**Figure S5: Distribution of Rho types in bacterial genomes (n = 2,730) associated with the phylogenetic tree based on the NCBI taxonomy.** Bacterial phyla, genome size (Mb), genome GC content (%), and sequence types are depicted. Rho domains were detected by hmmsearch using Pfam models and custom models built from Rho sequences deposited on the eggNOG v5 database. The phylogenetic tree was built with PhyloT v2 and annotated with iTOL v6.

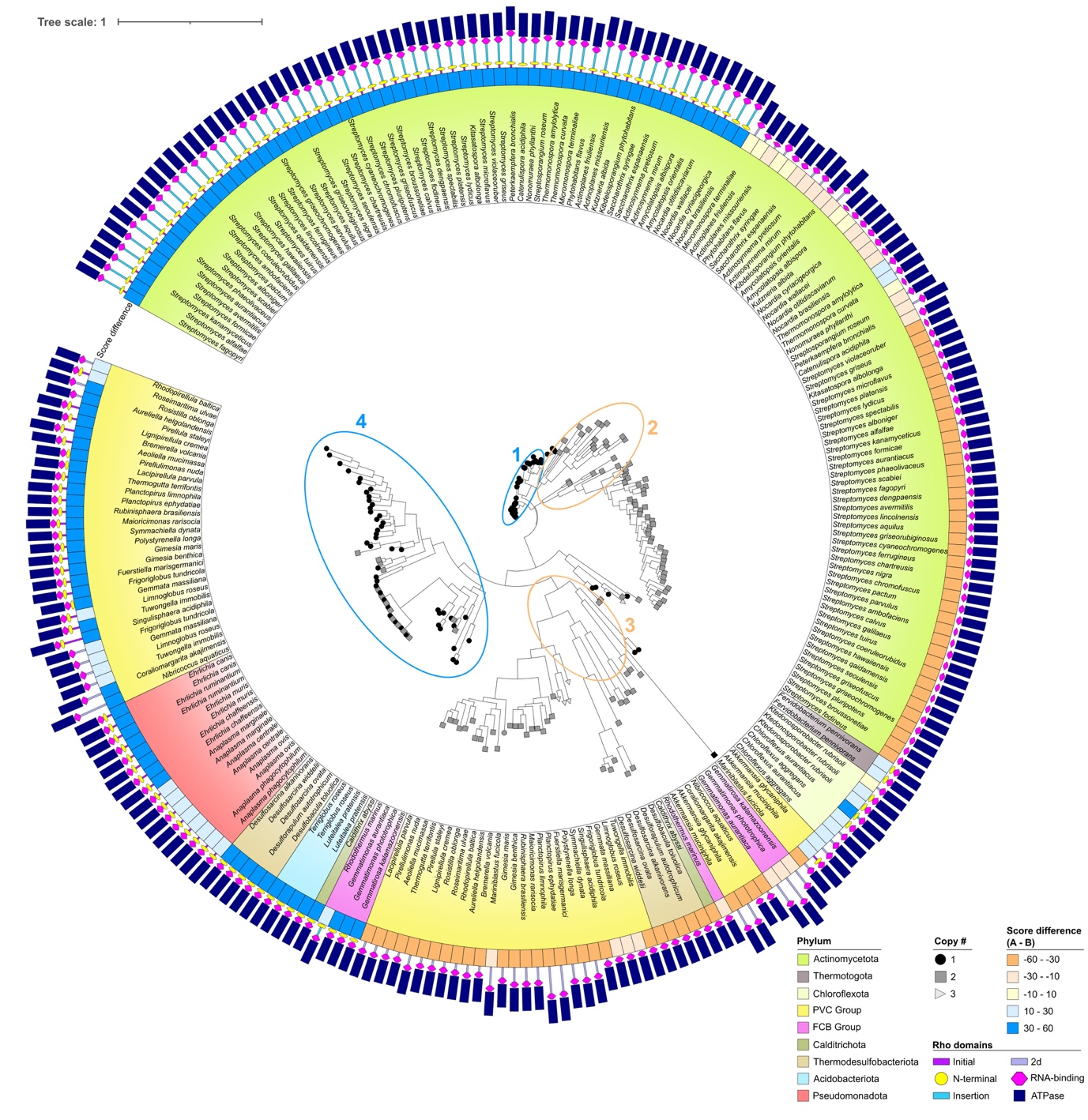

**Figure S6: Clustering of proteins from species with two Rho proteins.** The circular phylogram is based on the common regions corresponding to the RNA-binding + ATPase domains. Rho cluster into four distinct major groups with both the longer A (blue circles) and shorter B (orange circles) clustering into two major groups each. For each species (n = 116), the Rho copy number (sorted in the descending order of length) is represented by a geometric shape (circle: #1, square: #2, and triangle: #3) at the end of each tree node. The score difference between the A and B models is depicted with a color gradient from orange (higher score for B than A) to blue (higher score for A than B). The Rho sequences are represented by the Rho domains as geometric forms. The A and B models were developed by separating the two sequences of each species into groups of larger (A) and shorter (B) sequences (Methods). Sequence scores were calculated by hmmsearch using the developed models and the Rho duplicated sequences. The phylogenetic tree was built with FastTree after sequence alignment with MAFFT.

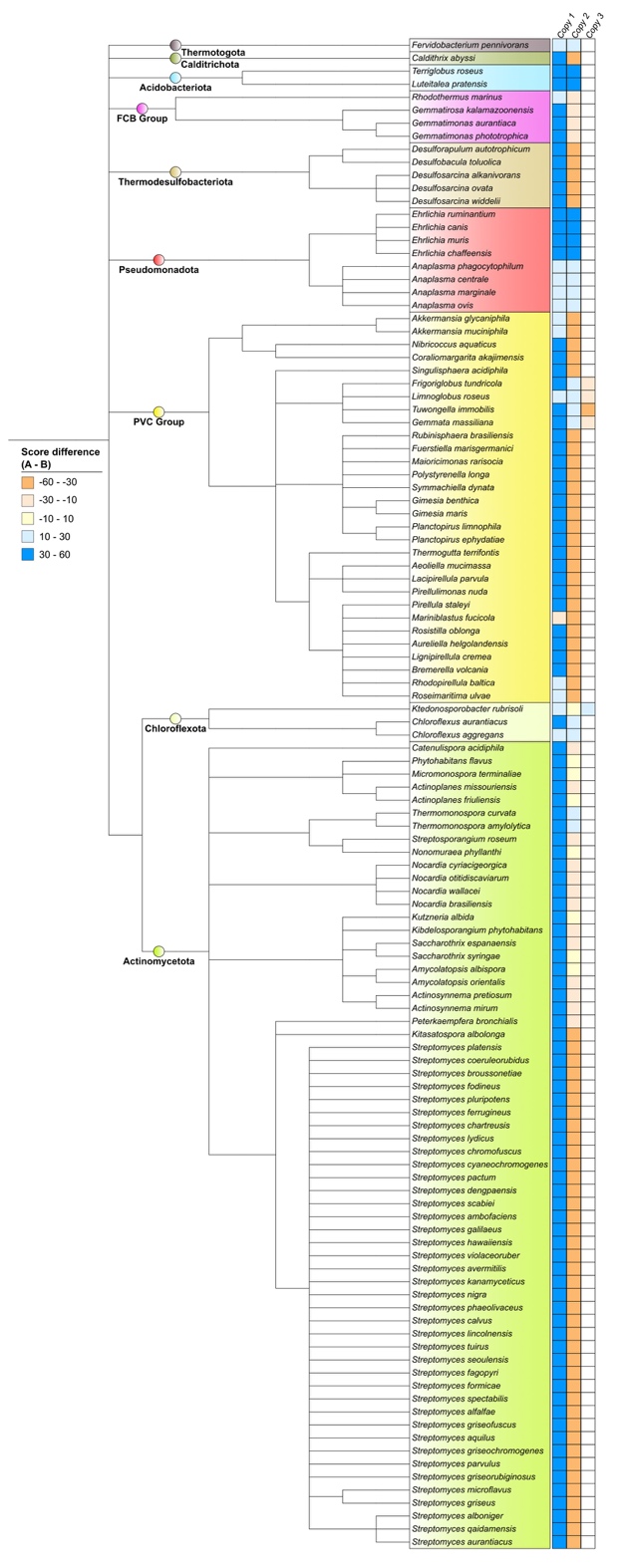

**Figure S7: Matches to A or B models in species with two or three copies of Rho.** The A and B models were developed by separating the duplicated sequences of each species into groups of larger (A) or truncated (B) sequences. Better matches to A (blue) or B (orange) models are shown as score differences (A - B) of Rho duplicated sequences associated with the phylogenetic tree based on the NCBI taxonomy. Most species have an A and a B, some species have two A like Rho (e.g. some Pseudomonadota) or B (*Mariniblastus fucicola*). Rho copy number corresponds to the sequence lengths sorted by descending order. Sequence scores were calculated by hmmsearch using the developed models and the Rho duplicated sequences. The phylogenetic tree was built with PhyloT v2 and annotated with iTOL v6.

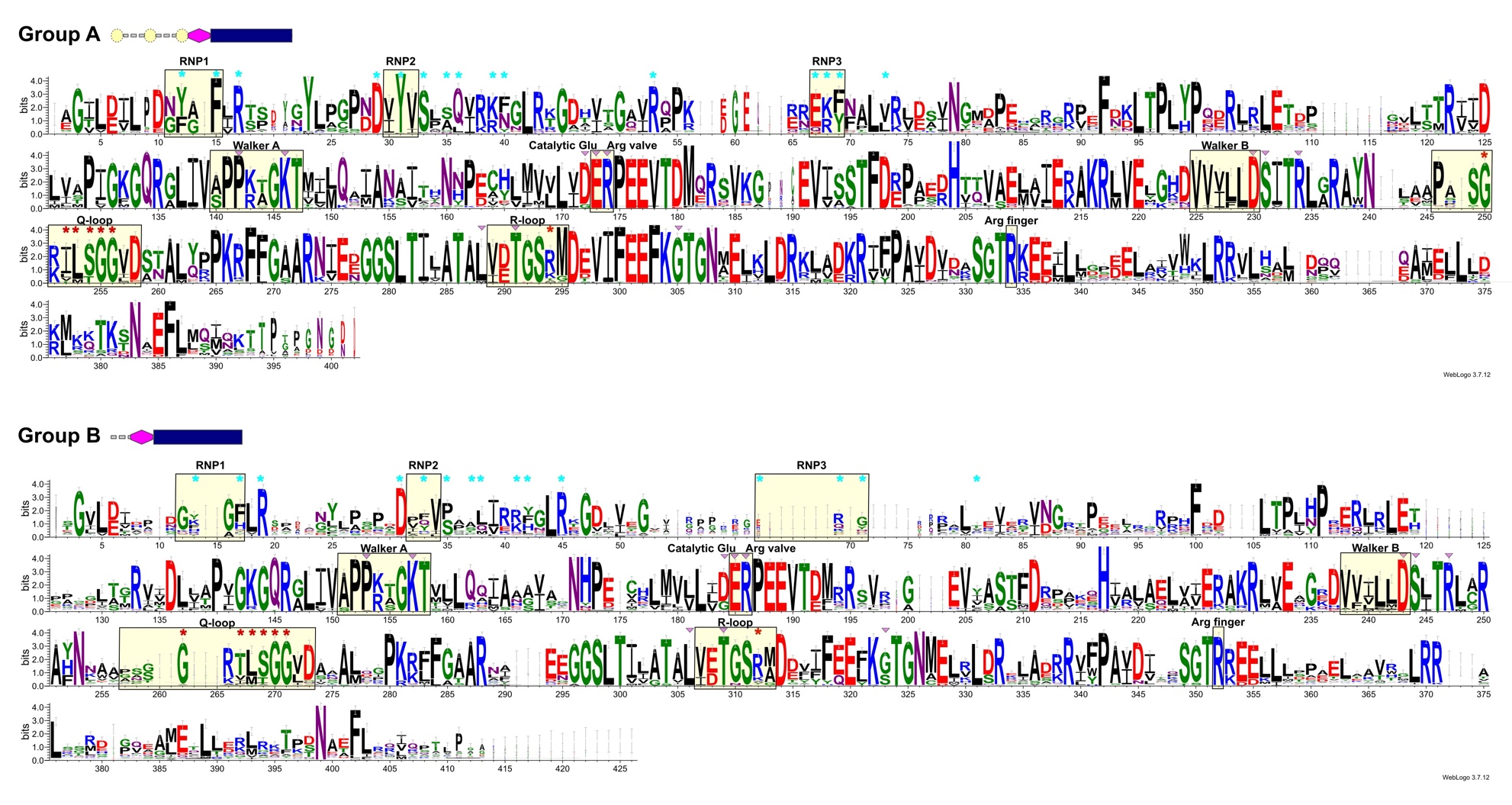

**Figure S8: Phylo-distinct Rho from species with two or more Rho.** (A) Alignment of the RNA-binding and ATPase regions of Group A Rho (mainly Type 2) (B) Alignment of the Group B Rho (mainly type 4). Group B have poorly conserved RNA binding domain motifs (RNP1, RNP2, RNP3). Blue stars indicate the positions of the primary RNA binding site in _Ec_Rho (PDB IDs: 1PVO and 8E6W). Pink triangles and red stars highlight the residues forming the BCM-binding pocket and the secondary binding site in _Ec_Rho (PDB ID: 1XPO), respectively. Sequence logos were generated from the sequence alignments on the WebLogo 3 server (https://weblogo.threeplusone.com).

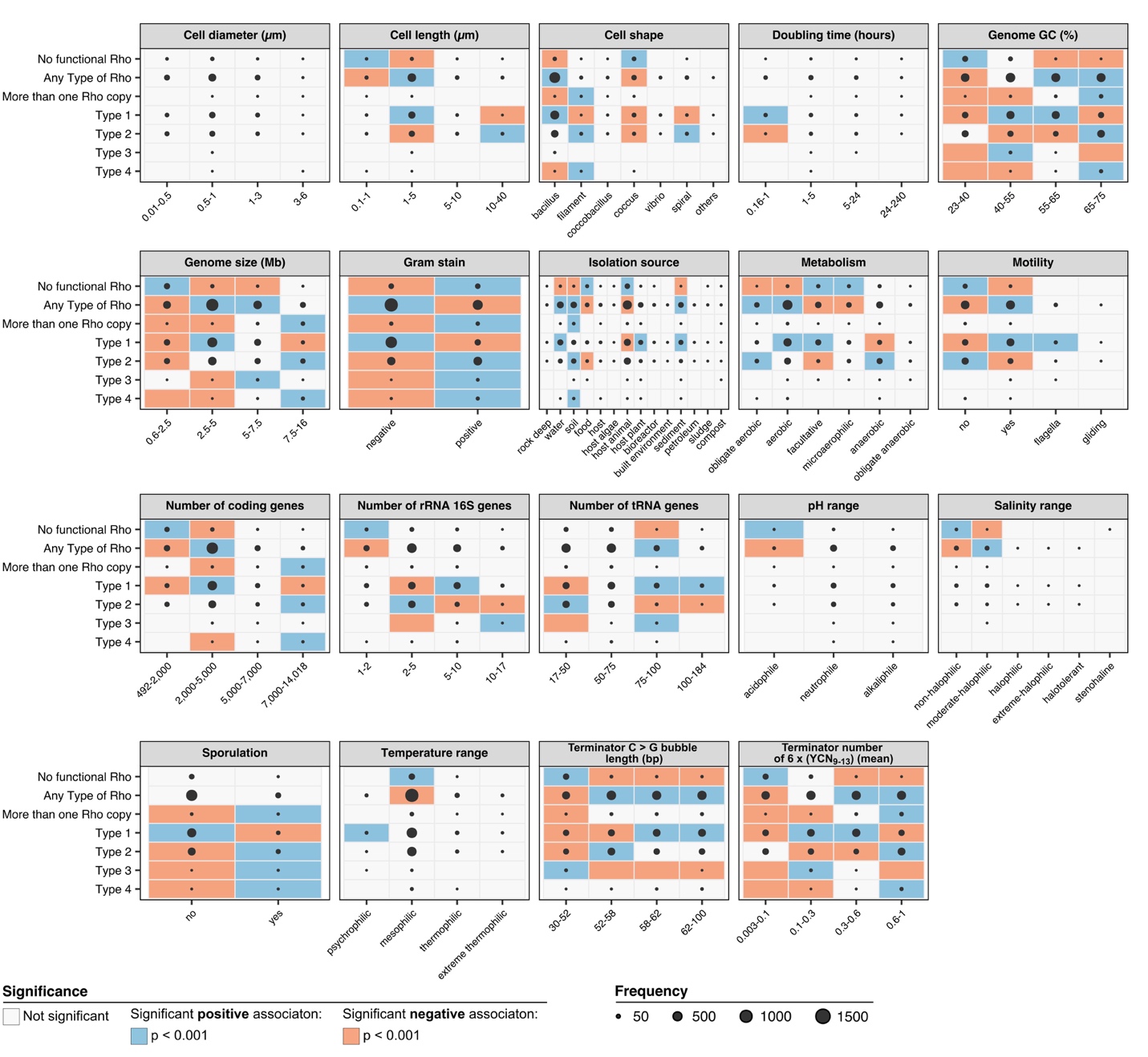

**Figure S9: Association of Rho types with bacterial genetic traits, habitats, and cell features.** Negative (orange) and positive (blue) associations between each type of Rho and each category were tested by one-sided Fisher’s exact test. The p-values were adjusted by Benjamini and Hochberg correction for multiple testing. The disk size represents the number of genomes (missing, if zero).

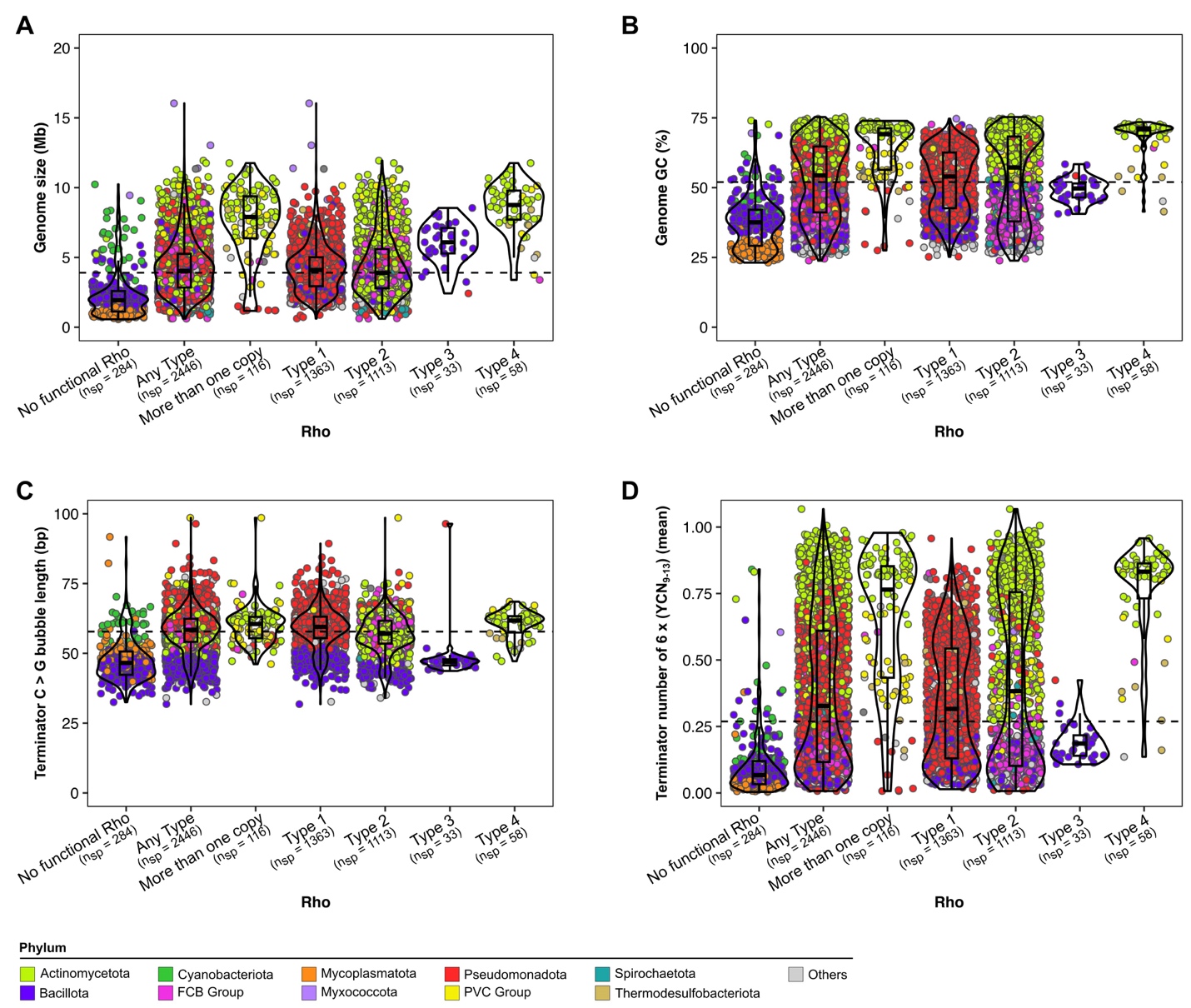

**Figure S10: The genomic features of bacterial species containing Rho groups indicating the bacterial phyla.** (A) Genome size (Mb). (B) Genome GC (%). (C) Mean C > G bubble length in terminators (bp) (D) Mean number of 6 x (YCN_9-13_) in terminators. The dashed line indicates the median values for genome size (3.86 Mb), GC% content (52%), mean C > G bubble length (bp) (57.8 bp), and number of 6 x (YCN_9-13_) in terminators (0.27). Box plots depict the distribution of the values as quartiles. The line at the center of each boxplot corresponds to the median value. Violin plots represent the density distribution of the values. The points are colored by the bacterial phyla.

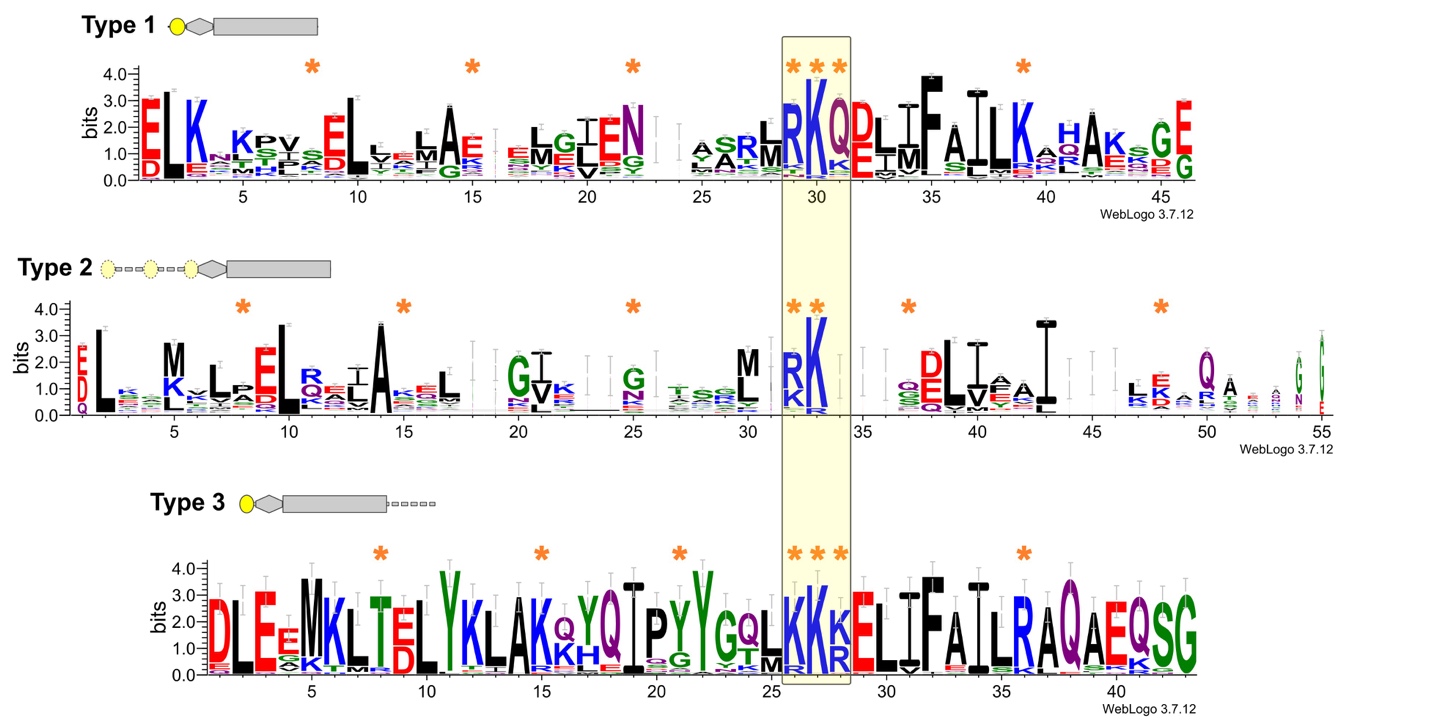

**Figure S11: Sequence logos of the N-terminal domain across the different groups of sequences harboring Rho domains.** Orange stars indicate the positions of the basic residues that create a positively charged patch in the N-terminal domain of Rho from *Thermotoga maritima* (PDB ID: 3L0O). Sequence logos were generated on the WebLogo 3 server (https://weblogo.threeplusone.com). * The N-terminal domain in the Type 2 sequences might be located at the beginning or after the initial region.

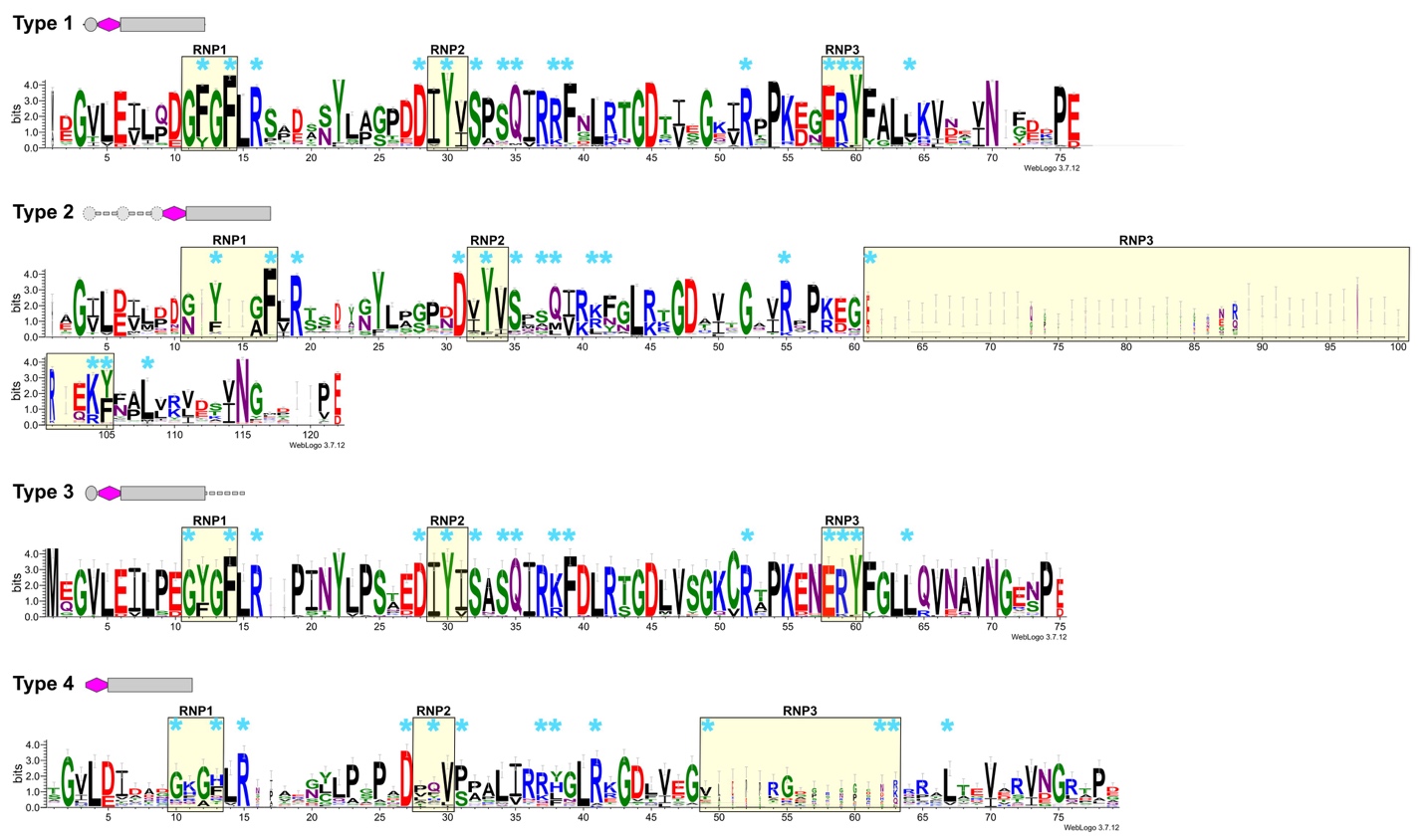

**Figure S12: Sequence logos of the Rho RNA-binding domain across the different groups of sequences harboring Rho domains.** Blue stars indicate the positions of the primary binding site in _Ec_Rho (PDB IDs: 1PVO and 8E6W). Sequence logos were generated on the WebLogo 3 server (https://weblogo.threeplusone.com).

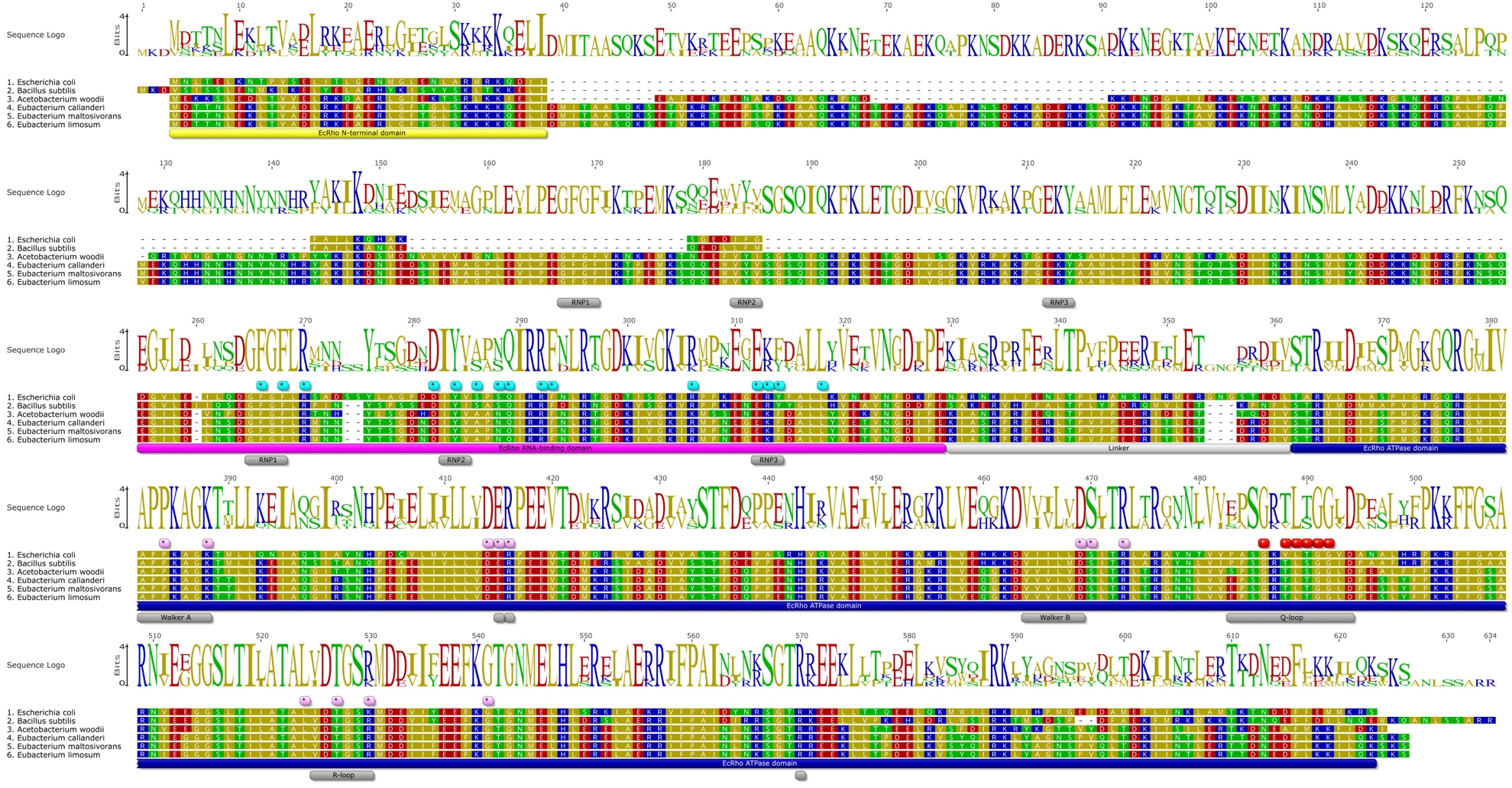

**Figure S13: Complete alignment of Rho sequences from species containing a double Rho RNA-binding domain.** The sequences of _Ec_Rho and _Bs_Rho (*Bacillus subtilis*) were used as reference. Blue stars indicate the positions of the primary binding site in _Ec_Rho (PDB IDs: 1PVO and 8E6W). Pink and red stars highlight the residues forming the BCM-binding pocket and the secondary binding site in _Ec_Rho (PDB ID: 1XPO), respectively. Alignments were performed by MUSCLE and colored according to the polarity scheme in Geneious Prime.

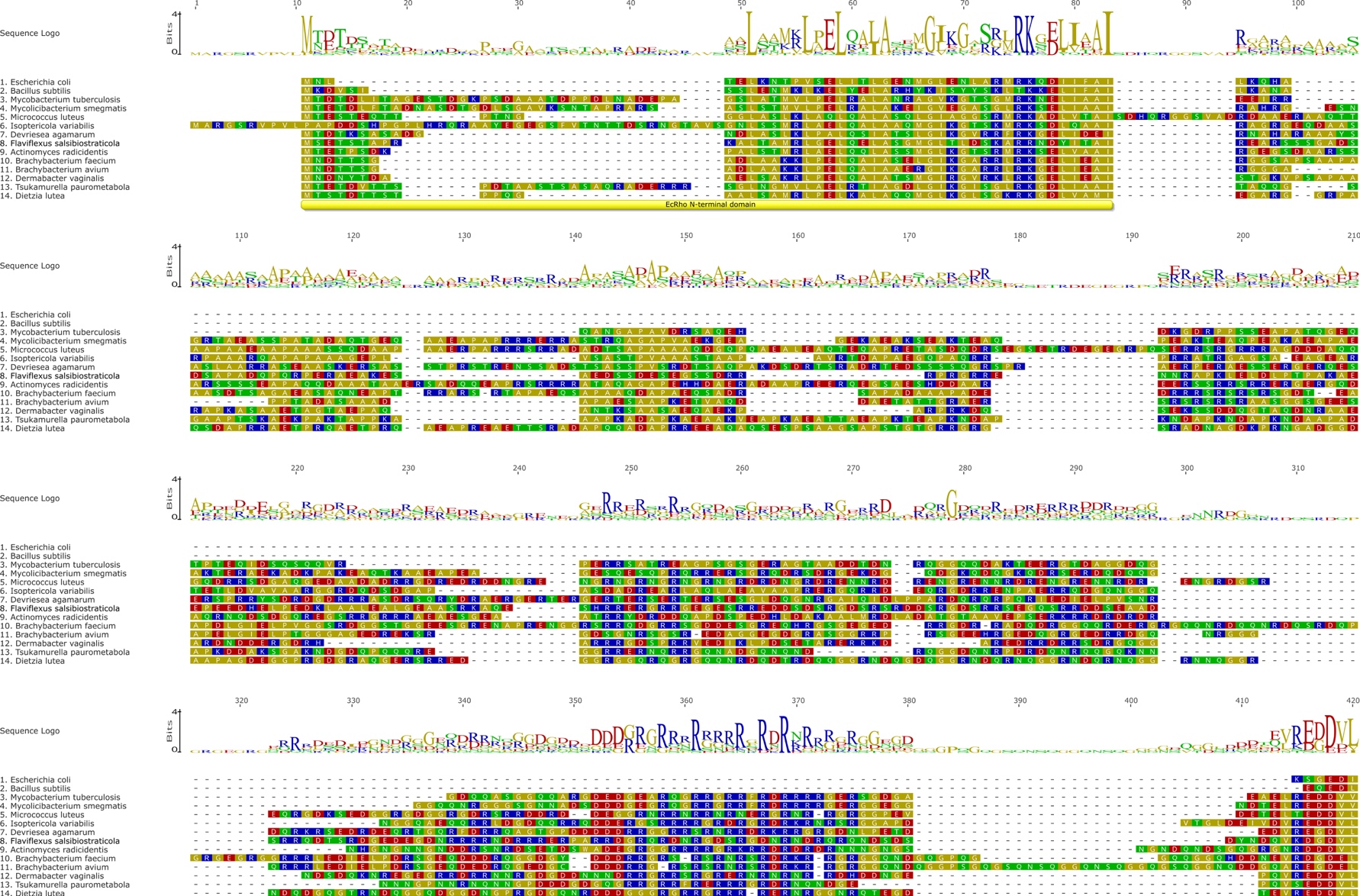

*(Continued on the next page)*

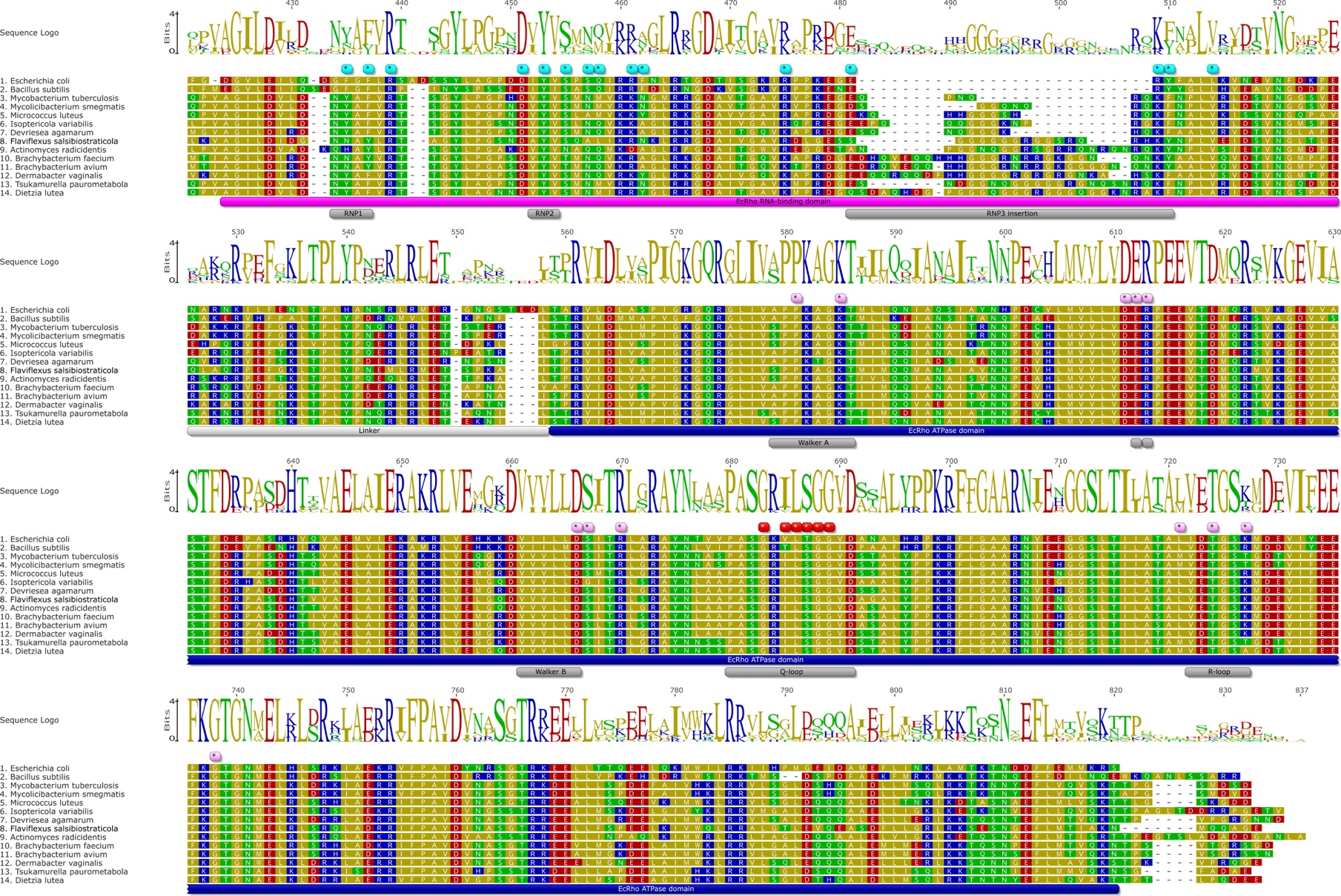

**Figure S14: Complete alignment of Rho sequences from species harboring the Rho RNA-binding domain with a RNP3 insertion.** The sequences of _Ec_Rho and _Bs_Rho (*Bacillus subtilis*) were used as reference. Blue stars indicate the positions of the primary binding site in _Ec_Rho (PDB IDs: 1PVO and 8E6W). Pink and red stars highlight the residues forming the BCM-binding pocket and the secondary binding site in _Ec_Rho (PDB ID: 1XPO), respectively. Alignments were performed by MUSCLE and colored according to the polarity scheme in Geneious Prime.

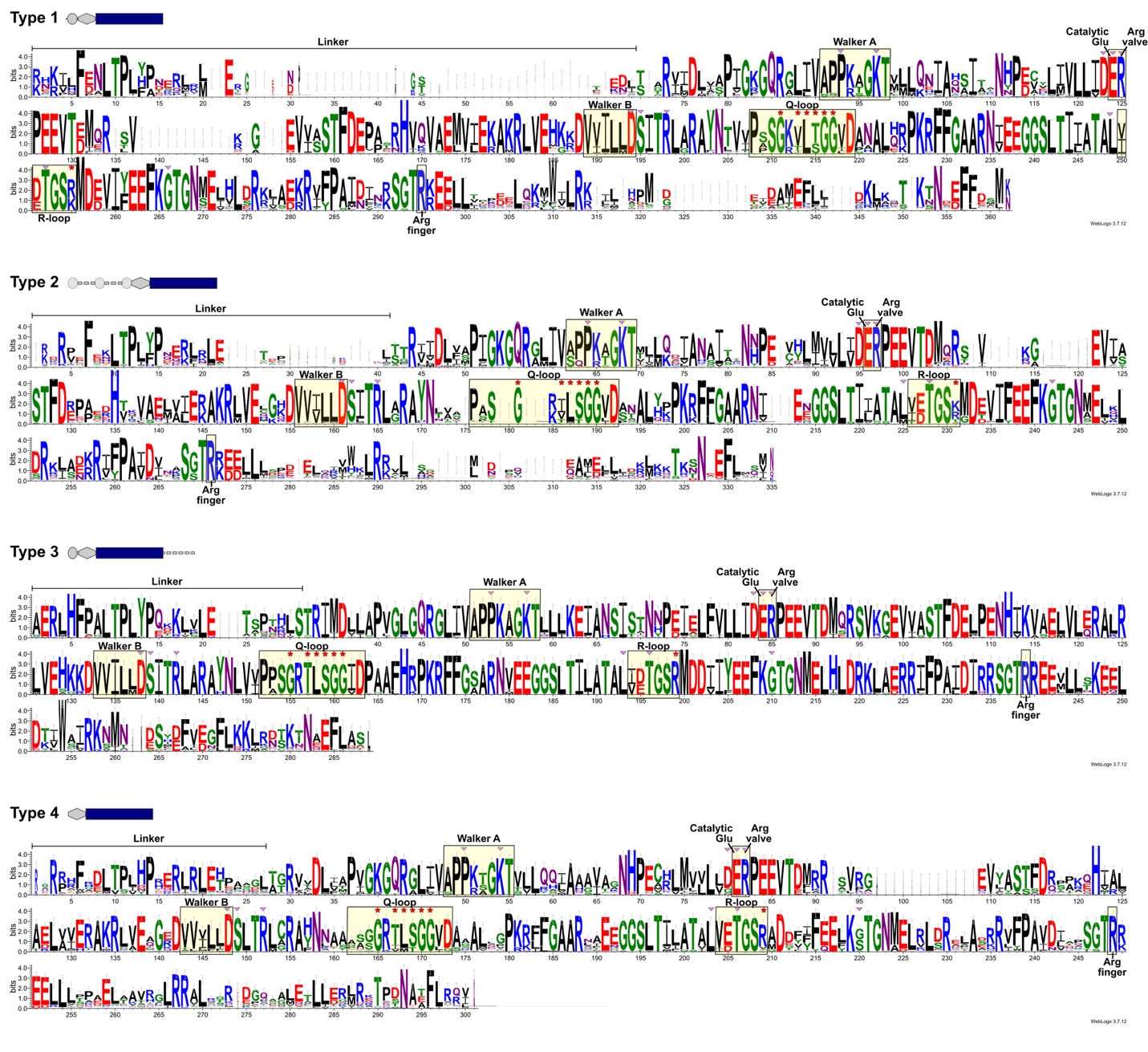

**Figure S15: Sequence logos of the Rho ATPase domain across the different groups of sequences harboring Rho domains.** Pink triangles and red stars highlight the residues forming the BCM-binding pocket and the secondary binding site in _Ec_Rho (PDB ID: 1XPO), respectively. Sequence logos were generated on the WebLogo 3 server (https://weblogo.threeplusone.com).

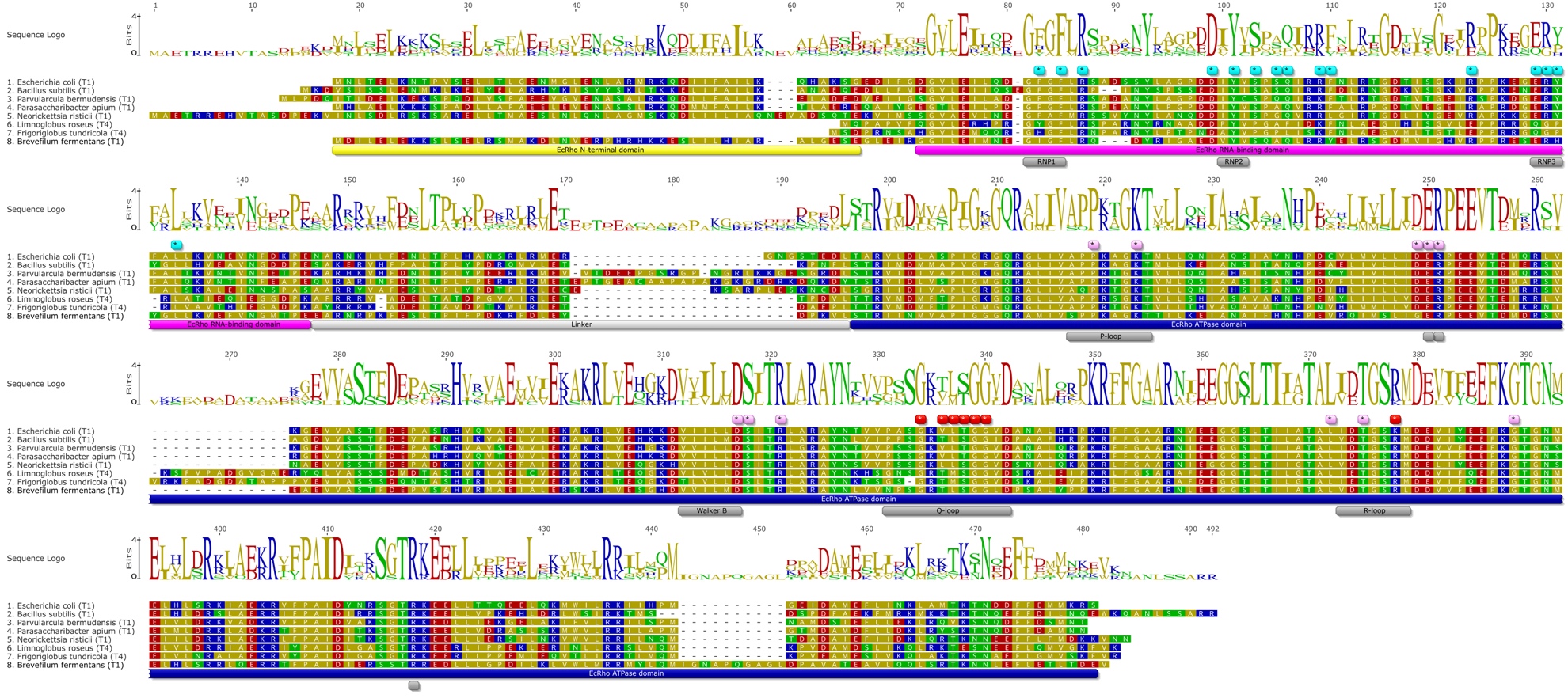

**Figure S16: Small insertions in the Rho ATPase domain.** The sequences of _Ec_Rho and _Bs_Rho (*Bacillus subtilis*) were used as reference. Blue stars indicate the positions of the primary binding site in _Ec_Rho (PDB IDs: 1PVO and 8E6W). Pink and red stars highlight the residues forming the BCM-binding pocket and the secondary binding site in _Ec_Rho (PDB ID: 1XPO), respectively. Alignments were performed by MUSCLE and colored according to the polarity scheme in Geneious Prime. All the ATPase domains with insertions are listed in Table S2 as are the available predicted AlphaFold structures.

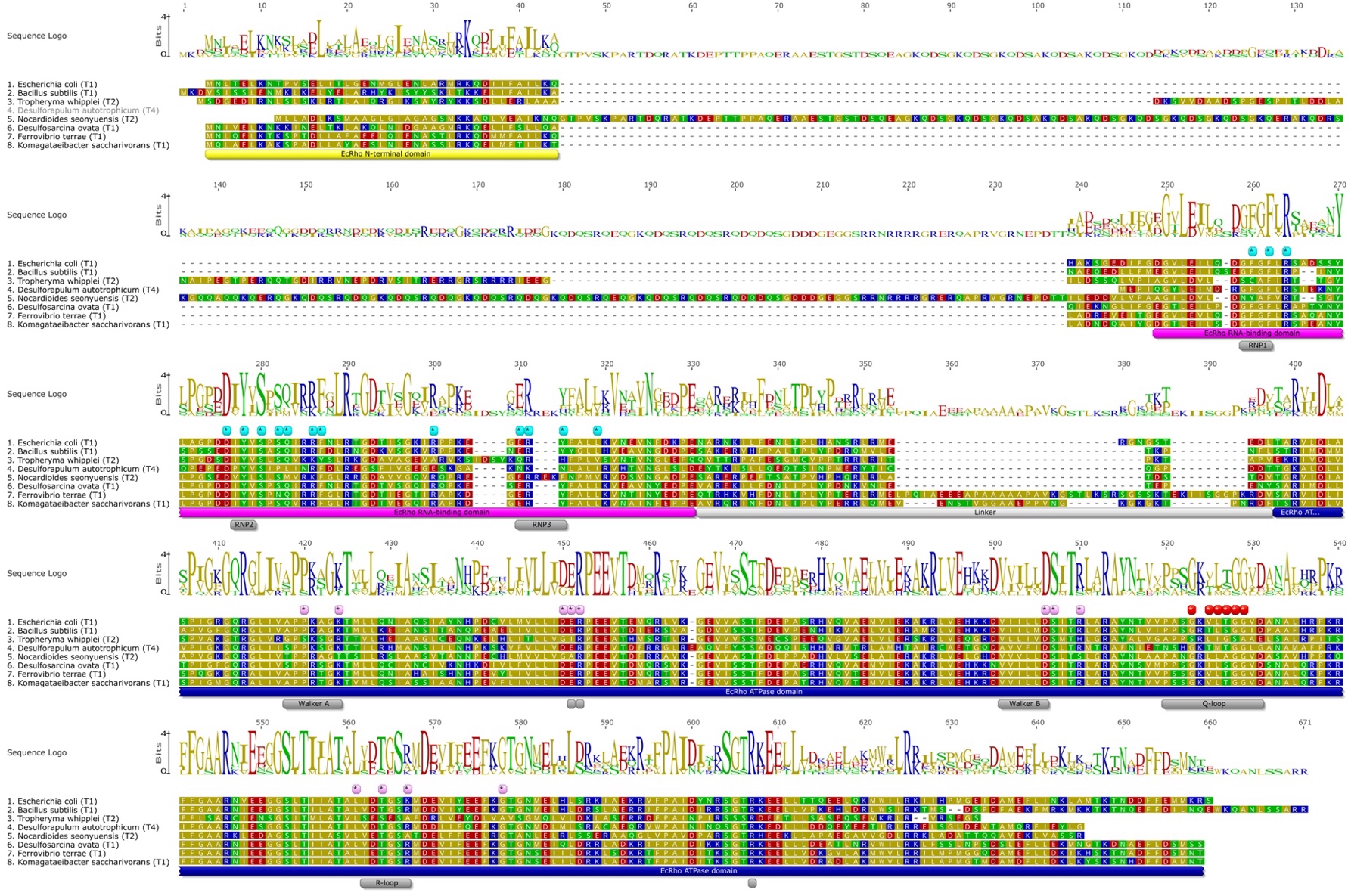

**Figure S17: Short deletions in the ATPase domain.** The sequences of _Ec_Rho and _Bs_Rho (*Bacillus subtilis*) were used as reference. Blue stars indicate the positions of the primary binding site in _Ec_Rho (PDB IDs: 1PVO and 8E6W). Pink and red stars highlight the residues forming the BCM-binding pocket and the secondary binding site in _Ec_Rho (PDB ID: 1XPO), respectively. Alignments were performed by MUSCLE and colored according to the polarity scheme in Geneious Prime. All the ATPase domains with deletions are listed in Table S2 as are the available predicted AlphaFold structures.

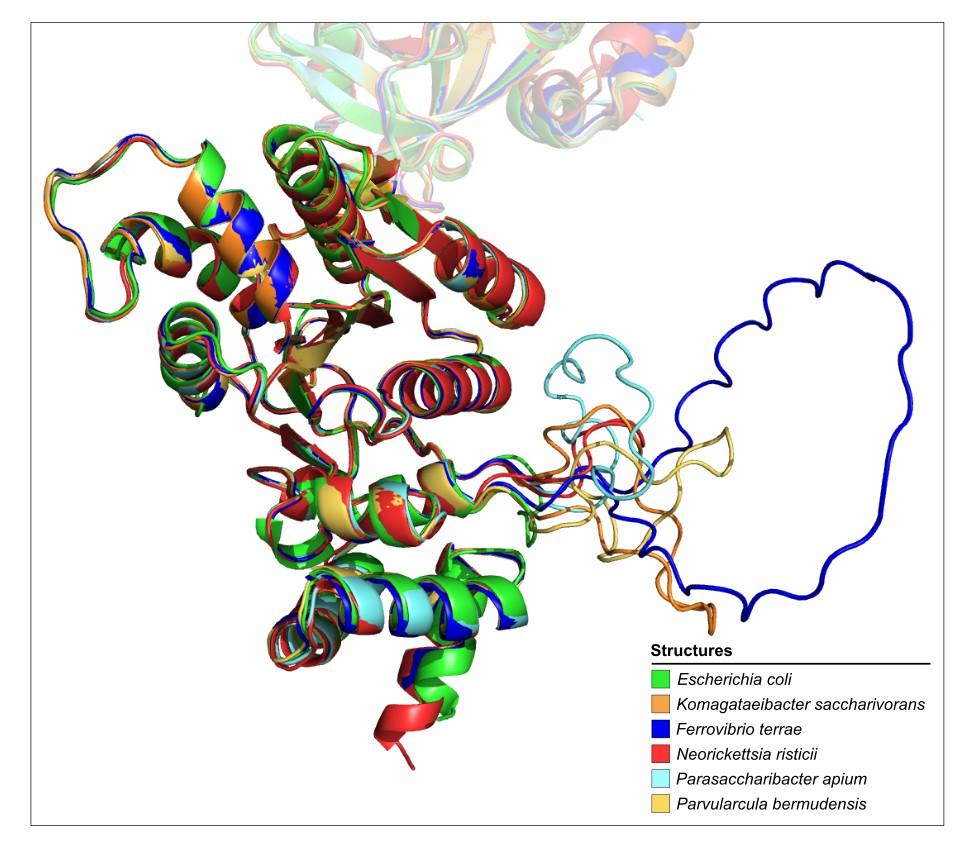

**Figure S18: AlphaFold predictions suggest that smaller insertions do not disrupt the overall Rho ATPase domain fold.** Rho secondary structures of *E. coli* (PDB ID: 8E6W), *Komagataeibacter saccharivorans* (AlphaFold ID: AF-A0A347WAH6-F1), *Ferrovibrio terrae* (AlphaFold ID: AF-A0A516GYM7-F1), *Neorickettsia risticii* (AlphaFold ID: AF-C6V3U9-F1), *Parasaccharibacter apium* (AlphaFold ID: AF-A0A7U7J0E3-F1), and *Parvularcula bermudensis* (AlphaFold ID: AF-E0TES2-F1).

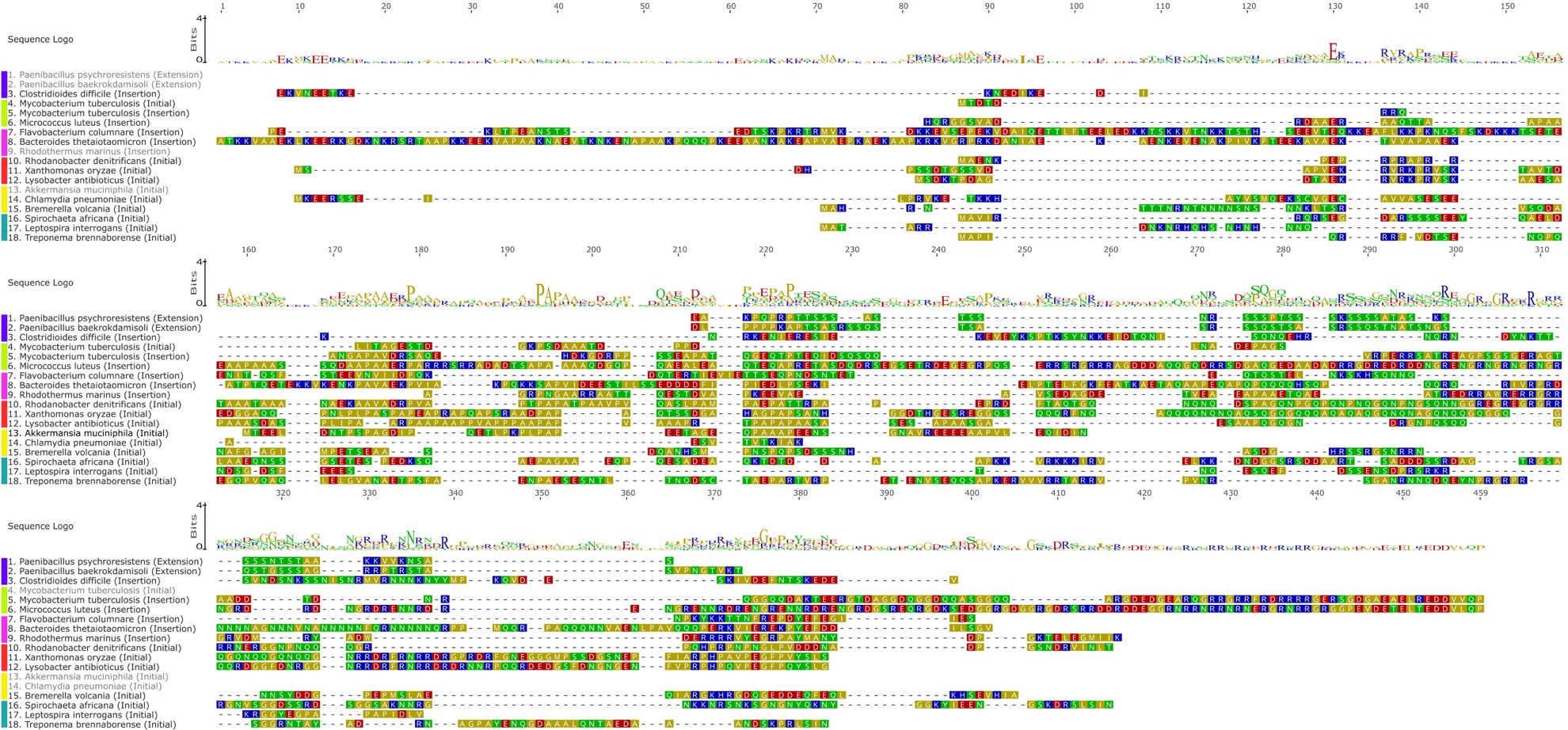

**Figure S19: A standard alignment of Rho additional regions does not identify sequence conservation**. Protein alignment of Rho extra regions (initial, insertion, and extension) from representative species of Bacillota (purple), Actinomycetota (light green), FCB Group (pink), Pseudomonadota (red), PVC Group (yellow), and Spirochaetota (green). The alignment was performed by MUSCLE and colored according to the polarity scheme in Geneious Prime.

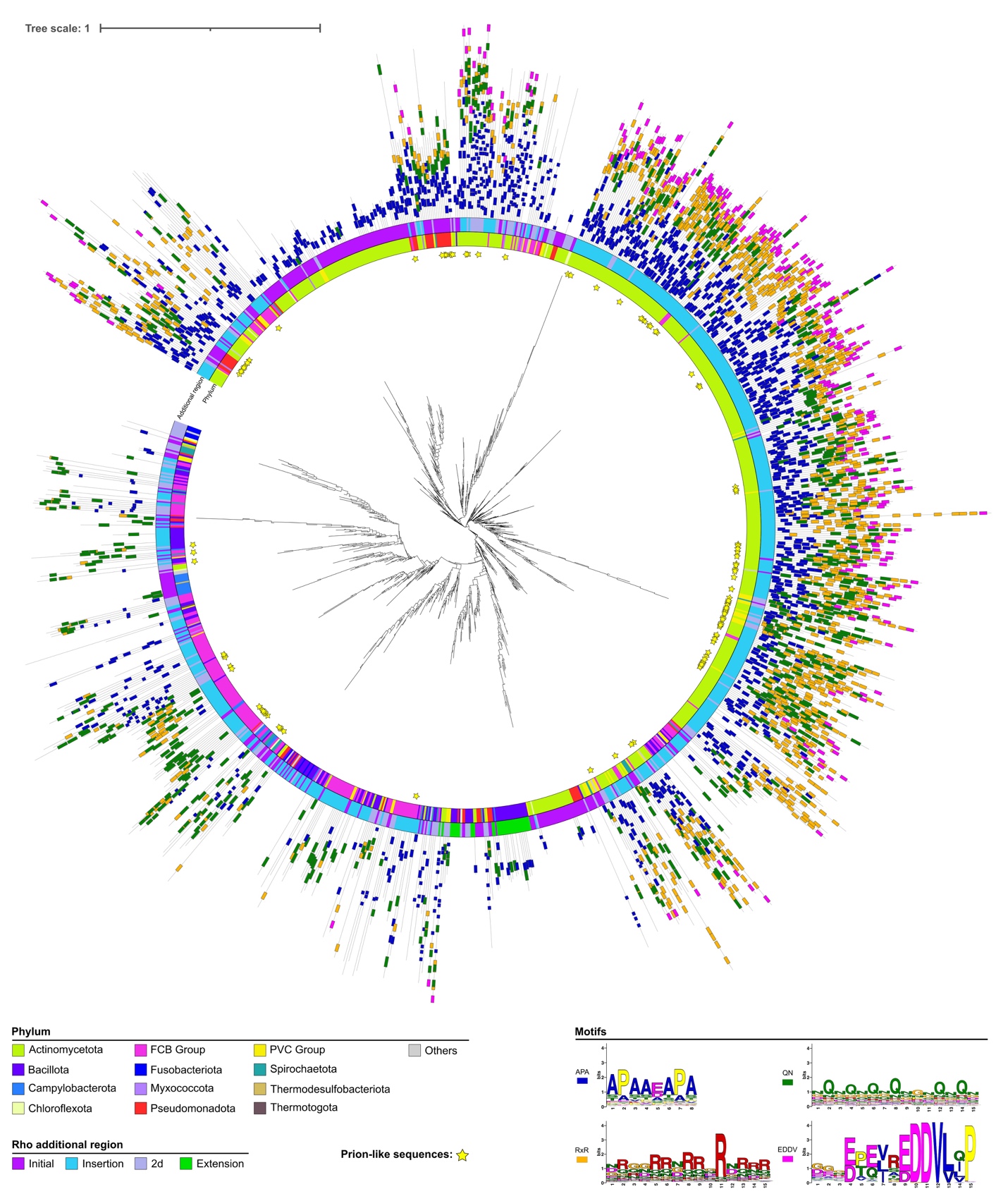

**Figure S20: Distribution and sequence logos of the significant motifs found in the additional regions of Rho factors.** Neighbor-Joining tree was constructed with MEGA after calculation of the distance matrix based on the alignment-free method. Bacterial phyla, Rho additional regions, Rho groups and significant motifs as represented in different colors. Sequence logos for the significant motifs were detected by MEME and confirmed with MAST in Rho additional regions.
